# Identification and characterization of BEND2 as a novel and key regulator of meiosis during mouse spermatogenesis

**DOI:** 10.1101/2021.11.05.467475

**Authors:** Longfei Ma, Dan Xie, Xiwen Lin, Hengyu Nie, Jian Chen, Chenxu Gao, Shuguang Duo, Chunsheng Han

**Affiliations:** State Key Laboratory of Stem Cell and Reproductive Biology, Institute of Zoology, Chinese Academy of Sciences, Beijing 100101, China; Innovation Academy for Stem Cell and Regeneration, Chinese Academy of Sciences, Beijing 100101, China; Institute of Zoology, University of Chinese Academy of Sciences, Chinese Academy of Sciences, Beijing 100049, China

## Abstract

The chromatin state undergoes global and dynamic changes during spermatogenesis, and is critical to chromosomal synapsis, meiotic recombination, and transcriptional regulation. However, the key regulators involved and the underlying molecular mechanisms remain poorly understood. Herein we report that mouse BEND2, one of the BEN-domain-containing proteins conserved in vertebrates, was specifically expressed in spermatogenic cells within a short time-window spanning meiotic initiation, and that it plays an essential role in the progression of prophase in meiosis I. *Bend2* gene knockout in male mice arrested meiosis at the transition from zygonema to pachynema, disrupted synapsis and DNA double-strand break repair, and induced non-homologous chromosomal pairing. BEND2 interacted with a number of chromatin-associated proteins—including ZMYM2, LSD1, CHD4, and ADNP— which are components of certain transcription-repressor complexes. BEND2-binding sites were identified in diverse chromatin states and enriched in simple sequence repeats. BEND2 contributed to shutting down the mitotic gene-expression program and to the activation of meiotic and post-meiotic gene expression, and it regulated chromatin accessibility as well as the modification of H3K4me3. Therefore, our study identified BEND2 as a novel and key regulator of meiosis, gene expression, and chromatin state during mouse spermatogenesis.

**Teaser:** Meiosis is a highly complex yet poorly understood process that involves the concerted actions of an increasing number of regulators, of which the list remains incomplete. Ma et al. identified BEND2 as a novel and key regulator of meiosis and showed that it interacts with critical chromatin modulators and specific genomic elements to control the expression of mitotic and meiotic genes.

## INTRODUCTION

Meiosis is the fundamental component of gametogenesis and consists of multiple processes that occur either sequentially or concurrently (*1, 2*). Meiosis is initiated when homologous chromosomes begin to pair and large-scale, programmed DNA double-strand breaks (DSBs) are generated (*3*). DSB repair and synapsis of homologous chromosomes are simultaneous and mutually dependent. Synapsis starts when the 3’ overhangs of DSBs invade homologous DNAs to form recombinant intermediates, and when the axial elements of the synaptonemal complex that consists of proteins such as SYCP3 and cohesin are bridged by SYCP1 and other central element components (*4*). Full synapsis is achieved when DSB repair intermediates are resolved into crossovers and the chromosomes become highly condensed around the complete synaptonemal complex.

These meiotic steps/processes are intricately coordinated by the complex interactions between chromatin and a large number of chromatin-binding proteins that include synaptonemal complex proteins, enzymes, chromatin modifiers/remodelers, and transcription factors (*5*). To give an example, PRDM9 acts as both a histone-modifying enzyme and a pioneer transcription factor, and interacts either directly or indirectly with many proteins—including CXXC finger protein 1 (CXXC1), EWS RNA binding-protein 1 (EWSR1), euchromatic histone lysine methyltransferase 2 (EHMT2), chromodomain Y like (CDYL), meiotic cohesin REC8, SYCP3, SYCP1, and lymphoid-specific helicase (LSH/HELLS) (*6, 7*). An increasing number of meiotic regulators have been identified by genetic studies, using model organisms such as gene knockout (KO) mice; these meiotic regulators include chromatin remodelers/modifiers and transcription factors such as HELLS (*8, 9*), YY1 (*10*), TET1 (*11*), INO80 (*12*), BRG1(*13*), Suv39h (*14*), DNMT3L (*15*), EHMT2 (*16*), PRDM9 (*17*), MLL2 (*18*), and SCML2 (*19*). Unfortunately, the spatiotemporal interactions between these regulators remain largely unknown. The concerted actions of these regulators usually result in chromatin states that are required for DNA activities such as DSB formation/repair, synapsis, and transcription (*20–24*). Specifically, correct chromatin states at particular genomic regions such as repetitive sequences and heterochromatin must be established to prevent erroneous recombination and/or transcription, which are detrimental to genome integrity (*10, 14*).

In the present study, we report the identification of a novel meiotic regulator, BEND2, that belongs to a BEN-domain-containing protein family that is poorly characterized. The BEN domain was first identified in diverse metazoan and viral proteins (usually with multiple copies), and was named after three experimentally characterized <u>p</u>roteins—BANP, <u>E</u>5R, and <u>N</u>AC1—in which it is present (*25*). A total of nine human and mouse genes that encode BEND1-9 are found in each of the genomes of these two species according to the NCBI Gene database. Although studies on the BEN family members are limited, they reveal the following key points: 1) BEN proteins tend to interact with a variety of proteins, most likely in a context-dependent manner; 2) most of the interacting proteins are components of transcription-repressive complexes involved in chromatin remodeling and/or modification; and 3) BEN proteins can be sequence-specific DNA-binding proteins (see Discussion for more details).

To our knowledge, there is no report regarding the functions of BEN proteins in germ cell development. In the present study, we showed that BEND2 is specifically expressed in spermatogenic cells shortly before and during prophase of meiosis I, and is essential for meiosis in male mice. We observed multiple meiotic defects in DSB repair and synapsis in male KO mice, such as complete spermatogenic arrest at zygonema. We also demonstrated that BEND2 interacts with multiple chromatin-binding proteins, and that it regulates chromatin states and transcription by preferentially targeting simple sequence repeats. These results add to our understanding of the molecular mechanisms governing meiosis and cell-specific regulation of chromatin states in meiotic cells.

## RESULTS

### BEND2 is a novel protein that is specifically expressed in spermatogenic cells around the time of meiotic initiation

We were initially interested in identifying and analyzing the functions of long noncoding RNA (lncRNA) genes that are specifically expressed in spermatogenic cells. An X chromosome-linked lncRNA gene based on the NCBI gene annotation was one such candidate, as we found that its transcripts were specifically expressed in mouse testes based upon our RNA-seq analyses of mouse multi-organ transcriptomic data (*26*). While our study was ongoing, this gene was re-annotated as encoding a protein belonging to the BEN family (*25*) (Fig. 1A). As the orthologous protein in humans has been named BEND2, we suggest that this mouse protein also adopt the same name. Predicted homologous BEND2 proteins can be found in vertebrates from fish to humans, and the sequence identity between the mouse and human proteins is 34% (Fig. S1A, B). The predicted longest transcript of mouse *Bend2* contains 15 exons, of which exons 2–15 harbor a coding sequence (CDS) for a protein of 728 aa (predicted molecular mass, 80 kDa) (Fig. 1A). And these transcripts were indeed detected exclusively in mouse testes by RT-PCR (Fig. 1B, S1C).

**Fig. 1.**
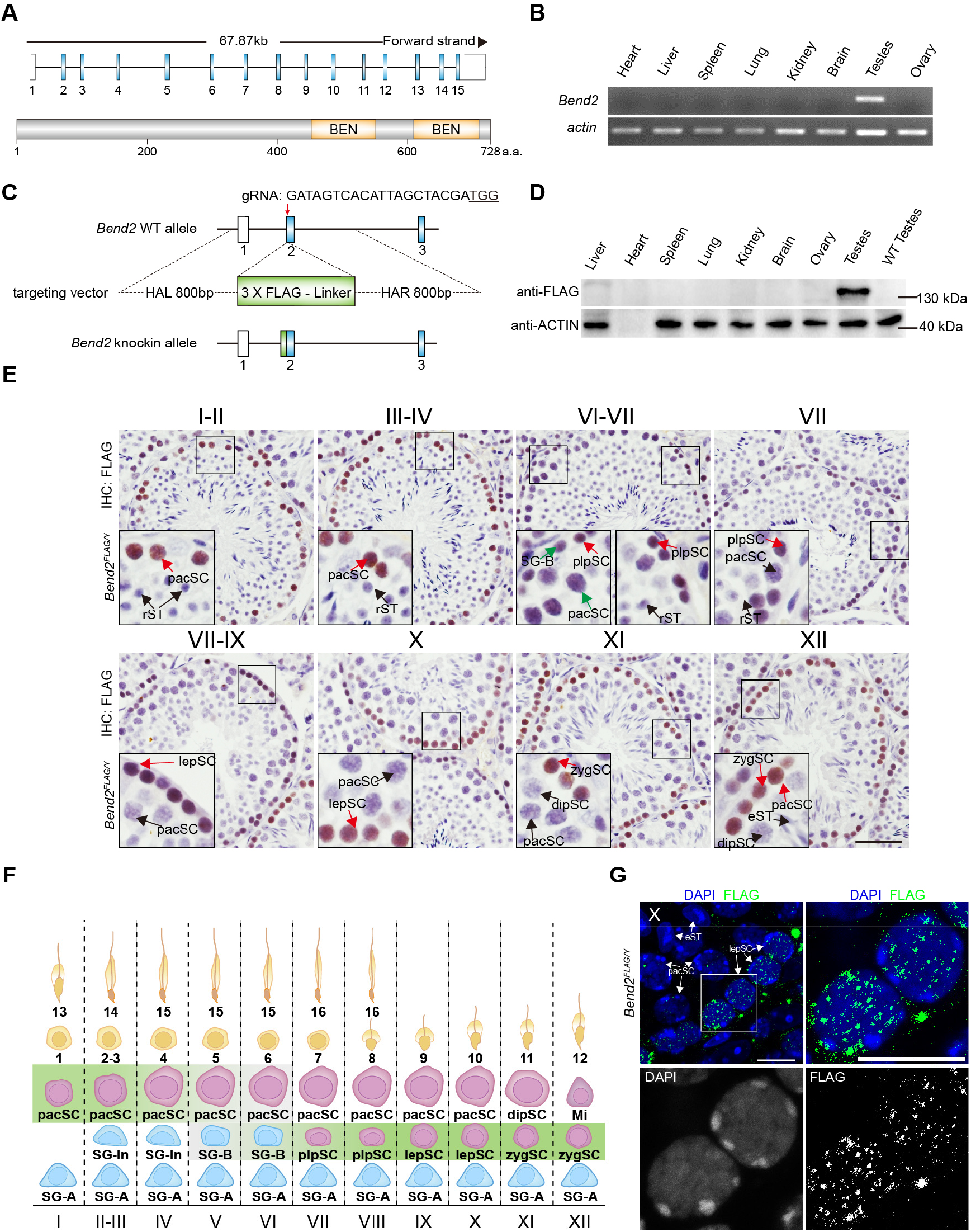
BEND2 is a novel protein specifically expressed around the time of meiotic initiation. **(A)** Schematic diagram of the primary structures of the *Bend2* gene and BEND2 protein. Upper diagram shows the structure of the *Bend2* gene; blue and white rectangles indicate protein-coded exons and UTR regions, respectively. The lower diagram represents the BEND2 protein, with orange boxes indicating the BEN domain. **(B)** RT-PCR detection of *Bend2* expression in multiple mouse organs. **(C)** Schematic representation of the locus of the 3 x FLAG tag knock-in; the tag sequence was inserted immediately behind the first codon of BEND2. **(D)** Western blotting analyses of BEND2 expression in multiple mouse organs using mmAb-FLAG. **(E)** Immunohistochemical staining of FLAG-BEND2 in testicular sections of various seminiferous stages using mmAb-FLAG; red and green arrows indicate BEND2-strongly- and -weakly-expressing cells, respectively; black arrows indicate no BEND2 expression. SG-B, type B spermatogonia; plpSC, pre-leptotene spermatocytes; lepSC, leptotene spermatocytes; zygSC, zygotene spermatids; pacSC, pachytene spermatocytes; dipSC, diplotene spermatocytes; rST, round spermatids; eST, elongated spermatids (scale bar, 20 μm). **(F)** Schematic summary of FLAG-BEND2 expression in male germ cell types and seminiferous stages. **(G)** Immunofluorescence staining of FLAG-BEND2 shown at higher magnification of BEND2 signals in leptotene spermatocytes (scale bar, 10 μm).

We developed a rabbit polyclonal antibody to BEND2 (rpAb-B2) by using a 30-aa synthetic polypeptide located between the two BEN domains (Fig. S1B). This antibody functioned appropriately in western immunoblotting and immunohistochemical analyses (Fig. S1D–F). By using rpAb-B2 in western blotting assays, we detected two proteins related to BEND2 in the testes of WT but not *Bend2* KO mice: one was 140 kDa (p140), while the other was 80 kDa (p80) (Fig. S1D). Intriguingly, p80 but not p140 could be consistently and specifically found in testes (Fig. S1E). As a non-specific protein was also detected by rpAb-B2 upon western blot analysis (labeled with an asterisk in Fig. S1D), we decided to use CRISPR-Cas9 technology to generate knock-in (KI) mice in which a 3XFLAG sequence was added to the N-terminus of BEND2 (FLAG-BEND2) (Fig. 1C, S1G). By using a mouse monoclonal Ab against FLAG (mmAb-FLAG), p140 but not p80 was detectable in the KI testes but not in the WT or the KO testes by westerns (Fig. S1H). p140 was also specifically observed when FLAG-BEND2 cDNA was expressed in 293FT cells by both rpAb-B2 and mmAb-FLAG, suggesting that p140 protein was FLAG-BEND2 itself (Fig. S1I, J). Based on these results, it is likely that p140 is the full-length BEND2, the mobility of which using SDS-PAGE was altered due to either post-translational modification(s) or unusual higher-order structures; p80 was a shorter version of BEND2, likely due to either alternative transcription or translation, or protein cleavage from p140. We next expressed the N- and C-terminal halves of BEND2 as FLAG-tagged proteins in 293FT cells (predicted molecular masses, 36 and 44 kDa, respectively), and found that the former migrated as a protein of 73 kDa while the latter migrated at 55 kDa. Sequence analyses showed that the N-terminal half of BEND2 was much more disordered and hydrophobic than the C-terminal half (Fig. S1K). Therefore, it was appropriate that BEND2 displayed slower electrophoretic mobility due to its usual sequence/structure at the N-terminus.

By using FLAG-BEND2 KI mice and the mmAb-FLAG antibody, we confirmed that p140 was exclusively expressed in testes among all of the tissues we examined (Fig. 1D). Using immunostaining of FLAG-BEND2 in testicular sections that could be staged based on hematoxylin staining, we found that the protein was highly expressed in preleptotene (plpSCs), leptotene (lepSCs), zygotene (zygSCs), and pachytene (pacSCs) spermatocytes from stages VII to III, and weakly in type B spermatogonia (SG-B) and pacSCs at stages V and VI (Fig. 1E, F). Interestingly, immunofluorescence imaging with higher magnification and a shorter exposure time revealed that the signals for FLAG-BEND2 in the nuclei of spermatocytes were punctate (Fig. 1G). These results indicated that BEND2 is an evolutionarily conserved novel protein, and that it is specifically expressed in the nuclei of spermatogenic cells in a stage-specific manner, shortly before and after meiotic initiation.

### *Bend2* gene knockout arrests spermatogenesis at prophase of meiosis I

We endeavored to assess the function of BEND2 by evaluating phenotypic changes in gene knockout (KO) mice that were generated by using CRISPR-Cas9 technology (*27*). Several founder mice with different mutant alleles were acquired, and a female (*Bend2*^-4k/+^) carrying a mutant allele with a 4-Kbp deletion (−4k) corresponding to a 104-aa deletion in the protein was crossed with WT males to at least the F2 generation for phenotypic evaluation (Fig. 2A–C, S2G). BEND2 was undetectable in the male founder with a 19-bp deletion (*Bend2*^-19/Y^) or male offspring from the founder with a single-base insertion (*Bend2*^+1/+^) (Fig. S2A, S2B).

**Fig. 2.**
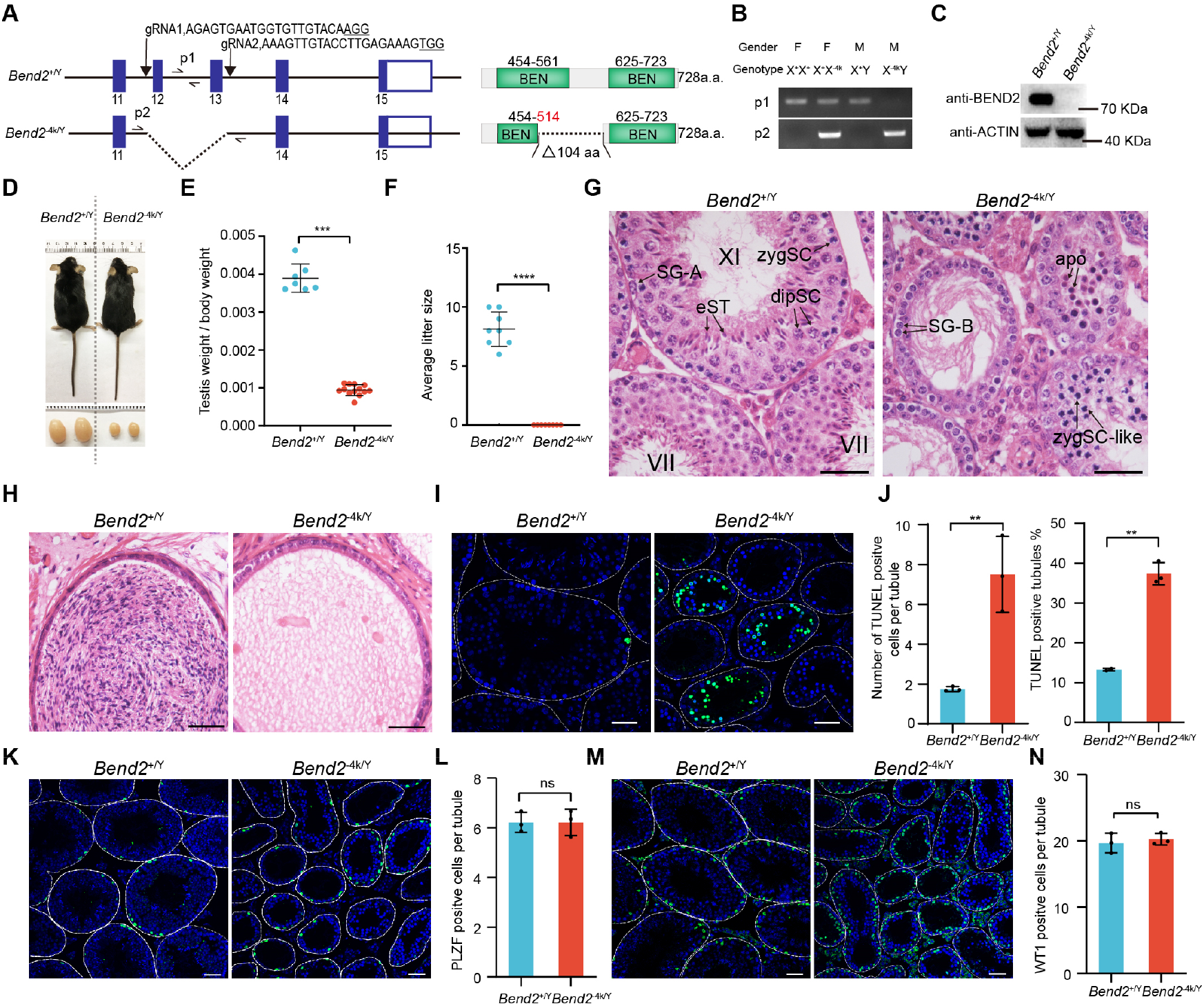
BEND2 is required for mouse spermatogenesis and the maintenance of male fertility. **(A)** Schematic illustration of the deletion of two exons of the *Bend2* gene to generate *Bend2*^-4k/Y^ mice; p1 and p2 indicate that two primers were used in mouse genetic identification. The right diagram shows BEND2 protein structure in WT and KO mice. **(B)** Identification of genotype with p1 (wild type allele) and p2 (mutant allele) primer pairs. **(C)** Western blot confirmation of the elimination of BEND2 protein in *Bend2*^-4k/Y^ mice using rpAb-B2. **(D)** Note the significant size reduction in 8-week-old *Bend2*^-4k/Y^ testes. **(E)** Quantitative comparison of testis/body ratios between *Bend2*^+/Y^ and *Bend2*^-4k/Y^ mice (\*\*\**p*<0.001, Student’s *t*-test). **(F)** Comparison of litter size in *Bend2*^+/Y^ and *Bend2*^-4k/Y^ mice (\*\*\*\**p*<0.0001). **(G)** H&E staining of testicular sections in *Bend2*^+/Y^ and *Bend2*^-4k/Y^ mice. zygSC-like, zygotene-like spermatocytes; apo, apoptotic cells (scale bar, 20 μm). **(H)** H&E staining of epididymal sections of *Bend2*^+/Y^ and *Bend2*^-4k/Y^ mice. **(I)** TUNEL staining of testicular sections in *Bend2*^+/Y^ and *Bend2*^-4k/Y^ mice; green signals indicate apoptotic cells (scale bar, 50 μm). **(J)** Quantitative comparison of TUNEL staining shows both TUNEL-positive cells and tubules were increased in *Bend2*^-4k/Y^ mice. **(K, L)** Immunofluorescence staining indicates that the number of PLZF-positive cells (green) were the same between *Bend2*^+/Y^ and *Bend2*^-4k/Y^ mice (scale bar, 20 μm). **(M, N)** Quantitative comparison of WT1 immunofluorescence staining of testicular sections in *Bend2*^+/Y^ and *Bend2*^-4k/Y^ mice (scale bar, 20 μm).

The *Bend2* KO male mice were infertile and exhibited markedly smaller testes than their wild-type littermates (Fig. 2D and E and Fig. S2C and D), and fertility testing showed that *Bend2* mutant males were infertile (Fig. 2F and Fig. S2E), and that they did not produce any haploid spermatids or spermatozoa (Fig. 2G, H, S2F). A closer inspection of the H&E-stained testicular sections revealed that the KO testes contained spermatogonia, leptotene and zygotene spermatocytes, and Sertoli cells— but no other type of germ cell (Fig. 2G). We also frequently observed apoptotic cells with condensed nuclei, and their presence was confirmed by TUNEL assays (Fig. 2G, I). Both the numbers of TUNEL-positive tubules and TUNEL-positive cells per tubule were significantly higher than numbers in the wild-type (WT) testes (Fig. 2J). In contrast, the numbers of undifferentiated spermatogonial stem cells (PLZF^+^) and Sertoli cells (WT1^+^) were equivalent between KO and WT mice (Fig. 2K–N). These results indicated that BEND2 plays a specific and essential role in meiosis in male mice.

### BEND2 occupies a role in DSB repair and synapsis

We next examined the molecular defects in the KO spermatocytes by immunostaining marker proteins involved in meiosis. The sub-stages of meiotic prophase I that include leptonema, zygonema, pachynema, and diplonema can be distinguished by the co-immunostaining patterns of SYCP3 and the phosphorylated form of histone H2AX (γH2AX) that marks DSBs formed during meiosis. Under normal conditions, γH2AX signals in lepSCs and zygSCs are diffusely localized over the nuclei, indicating large numbers of unrepaired DSBs. In contrast, in pacSCs and dipSCs, the signal is a small dot that marks a territory occupied by the X and Y chromosomes (also known as the sex body). The clearance of γH2AX from the nuclei of pacSCs (except for the sex bodies) indicates that DSBs in the autosomes have been repaired. pacSCs were easily identified in WT testes by the co-staining of γH2AX and SYCP3, but they were absent in the KO testes (Fig. 3A). As some tubules contained only dot-shaped or diffuse γH2AX signal while others contain both, three types of tubules (dot-only, diffusion-only, and double-stained) and two types of cells (dot and diffusion) were observed in WT testes (Fig. 3B, C, S3A). In contrast, only diffusion-only tubules and diffusion cells were seen in KO testes. Moreover, the numbers of both diffusion-only tubules and diffusion cells were much higher in KO testes than in WT testes (Fig. 3B, C, S3A). These results indicated that DNA DSB were formed but not properly repaired in *Bend2* KO mice.

**Fig. 3.**
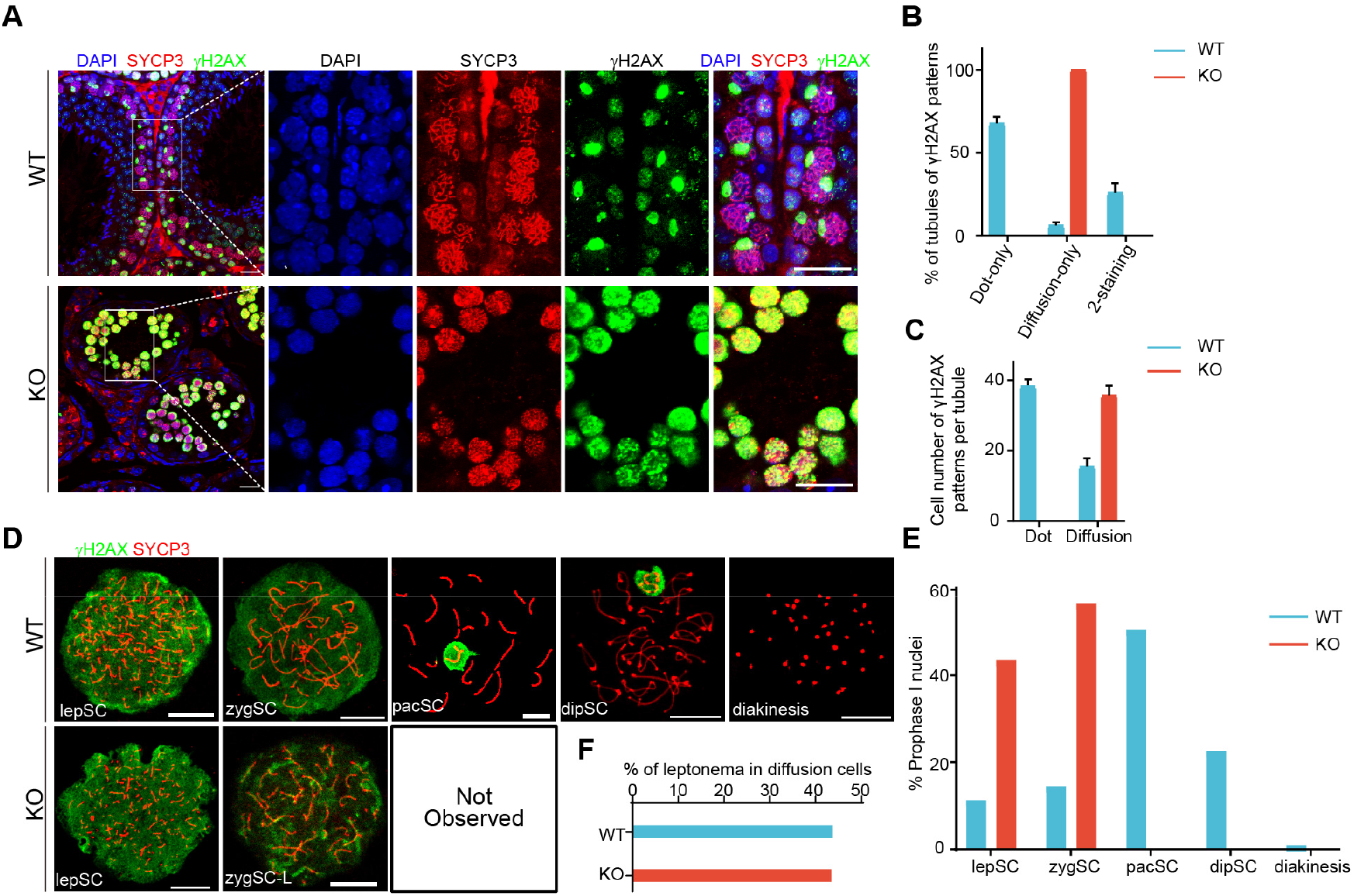
Spermatocytes with *Bend2* knockout fail to complete meiotic prophase. **(A)** Immunofluorescent labeling of testicular sections with mouse polyclonal SYCP3 antibodies (red) and rat polyclonal γH2AX antibodies (green). DNA was counterstained with DAPI (blue) and merged images are shown (scale bar, 10 μm). **(B)** Proportion of tubules with γH2AX expression patterns. Tubules of each mouse were counted: for *Bend2*^+/Y^, n= 354; for *Bend2^-^*^4k/Y^, n=321. **(C)** Average number of spermatocytes with γH2AX expression patterns per tubule. At least 100 tubules of each mouse were counted. **(D)** Nuclear spreads of various spermatocytes in *Bend2*^+/Y^ and *Bend2^-^*^4k/Y^ mice. Spermatocytes were immunostained with SYCP3 (red) and γH2AX (green) (scale bar, 10 μm). **(E)** Frequency statistics for spermatocytes in the meiotic-prophase I stage for WT and KO mice. Number of spermatocytes analyzed: for *Bend2*^+/Y^, n= 256; for *Bend2^-^*^4k/Y^, n=180. **(F)** Proportions of lepSC in γH2AX diffusion cells.

Co-immunostaining of SYCP3 and γH2AX was also carried out on surface-spread spermatocytes to reveal details that were invisible in testicular sections (Fig. 3D and Fig. S3B). While all spermatocytes from leptonema to diakinesis of meiosis I were observed in WT testes, pacSCs with dot-shaped γH2AX signals and subsequent cell types were never observed in KO mice (Fig. 3D, Fig. S3B). Moreover, the SYCP3-labeled chromosomal axes in KO zygSCs were not typical of the long continuous threads in WT zygSCs; rather, the former were more condensed, and we therefore named them zygSC-like cells (zygSC-LCs). Quantitative analyses showed that the percentages of lepSCs and zygSC-LCs among all cells at prophase of meiosis I were much higher in KO testes (Fig. 3E). As the diminution in the total number of spermatocytes contributed to the increase in the percentages of lepSC and zygSC-LCs in KO testes, when we calculated the percentages of lepSCs of cells with diffuse γH2AX signals, we found that the percentage did not change between KO and WT mice (Fig. 3F). This suggested that the absolute number of lepSCs was increased commensurately in KO mice as the total number of cells with diffuse γH2AX staining per tubule was increased.

The progression of synapsis between homologous chromosomes can be monitored by the co-staining of SYCP3 and SYCP1. Under normal conditions, SYCP3 but not SYCP1 can be detected in lepSCs; SYCP1 is initially detectable in zygSCs as short segments along the relatively more continuous SYCP3 threads; and it then becomes fully co-localized with SYCP3 to form the thick, smooth, and individualized synaptonemal complex in pacSCs. Surprisingly, we noted SYCP1 in approximately 29% of KO lepSCs (Fig. 4A, S4A), and we identified three types of zygSC-LCs (zygSC-L1, 46%; zygSC-L2, 28%; zygSC-L3, 26%) in KO testes (Fig. 4B). SYCP1 signals were primarily observed as small dots in zygSC-L1, while in zygSC-L2, they were thick or thin segments that represented the synaptonemal complex between homologous chromosomes and sister chromatids, respectively. In zygSC-L3, the SYCP1 signal was mostly detected as 40 discontinuous thin segments representing 40 univalents that underwent synapsis between sister chromatids.

**Fig. 4.**
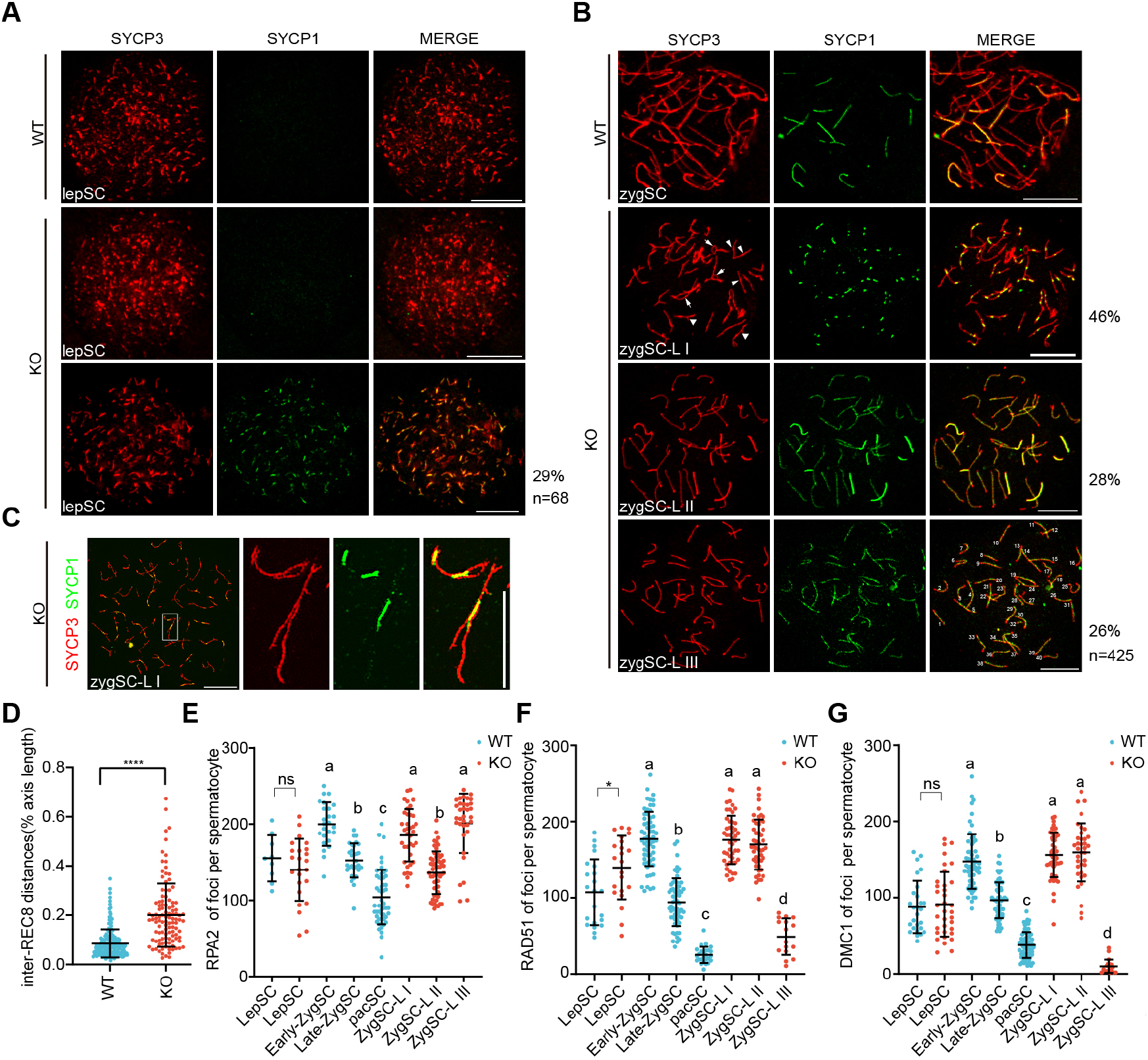
BEND2 is required for homologous synapsis in meiosis. **(A-B)** Immunofluorescent labeling of SYCP3 (red) and the transverse filament protein SYCP1 (green), a marker of synapsis. **(A)** Abnormal SYCP1 signals were observed in leptotene spermatocytes of *Bend2*^-4k/Y^ mice compared with *Bend2*^+/Y^ mice. Approximately 30% of leptotene spermatocytes were abnormal in *Bend2*^-4k/Y^ mice (n=68). **(B)** SYCP3- and SYCP1-staining of *Bend2^+/Y^* zygotene spermatocytes and *Bend2*^-4k/Y^ zygotene-like spermatocytes. According to their staining with SYCP1 and SYCP3, zygotene-like spermatocytes were divided into three classes: zygSC-L I, 46%; zygSC-L II, 28%; and zygSC-L III, 26% (n=425). **(C)** Super-resolution microscopic images of zygotene-like spermatocytes showing abnormal synapsis (scale bar, 10 μm). **(D)** Scatterplot in which we compared inter-REC8 distances along chromosomes in *Bend2*^+/Y^ and *Bend2*^-4k/Y^ mice (*p*<0.0001, obtained with two-tailed, unpaired *t*-test). **(E)** Each dot represents the number of RPA2 foci per spermatocyte; solid lines show the mean and SD of focus number in each group of spermatocytes. Data were analyzed with one-way ANOVA and Tukey’s multiple-comparison test. **(F)** Each dot represents the number of RAD51 foci per spermatocyte. Solid lines show the mean and SD of focus number in each group of spermatocytes. Data were analyzed with one-way ANOVA and Tukey’s multiple comparison test. **(G)** Each dot represents the number of DMC1 foci per spermatocyte. Solid lines show the mean and SD of focus number in each group of spermatocytes. Data were analyzed with one-way ANOVA followed by Tukey’s multiple comparison test.

As inter-sister synapsis was first uncovered in mice with cohesin gene *Rec8* knockout and that was also observed in KO/mutant mice of several other cohesin genes, we examined whether REC8 foci were modified in *Bend2* KO mice (*28*). Because the REC8 foci were numerous and not well separated from each other, we measured the distances between well-separated foci along chromosomal axes in zygSCs and zygSC-LCs, and found that the average distance in KO mice was significantly longer than in WT littermates, suggesting a reduced number of REC8 foci in zygSC-LCs (Fig. 4D, S4C). Notably, SYCP3 signals in the form of forks, bubbles, and unequal branches were frequently detected in zygSC-L1 (Fig. 4B). The presence of multiple and unequal branches was more evident in images from super-resolution structured illumination microscopy (SIM), and signified synapses between nonhomologous chromosomes (Fig. 4C, S4B). These results suggested that synapses initiated prematurely in KO mice (as early as in lepSCs), but could not be fully established between homologs; instead, they progressed in incorrect directions to form non-homologous and inter-sister synapses in different zygSC-LCs.

Meiotic DNA DSBs are repaired in a step-wise manner. The DNA ends of DSBs are resected into long single-stranded 3’ overhangs that are initially coated by RPA proteins. RPAs are subsequently replaced by the recombinase proteins RAD51 and DMC1 to form nucleoprotein filaments that seek a homologous template and form the recombinant intermediate, and are finally resolved into either crossovers between homologs or non-crossovers (*29, 30*). We distinguished the DNA-bound RPA2, RAD51, and DMC1 as hundreds of foci along the SYCP3 threads in lepSCs and zygSCs by co-immunostaining (Fig. S4E–G). In WT mice, the focus numbers of all three proteins increased from leptonema to early zygonema, decreased from early to late zygonema, and reached their nadirs at pachynema (Fig. 4E–G). In lepSCs, both the numbers of foci for RPA2 and DMC1 were similar between WT and KO mice, while the number of RAD51 foci in KO mice was higher than in WT controls. The numbers of RPA2 foci in zygSC-L1 and L2 were similar to those in early and late zygSCs in WT mice, respectively. Of note, the number of RPA2 foci in zygSC-L3 was also similar to that in early zygSCs. The numbers of both RAD51 and DMC1 foci were similar among early zygSCs, zygSC-L1 and zygSC-L2. Of greater interest, the number of RAD51 foci in zygSC-L3s was lower than in late zygSCs but higher than pacSCs while the number of DMC1 foci in zygSC-L3 was the lowest among all cell types. These results suggested the following: 1) that more recombinant intermediates were formed in KO lepSCs than in WT ones, consistent with the appearance of a significant proportion of SYCP1^+^ lepSCs in KO mice; 2) that the formation of recombinant intermediates in zygSC-L1 and zygSC-L2 was relatively normal compared with that in early zygSCs of WT mice, suggesting these two KO cell types are still at the early zygonema with aberrant synapses; 3) that zygSC-L3 underwent inter-sister synapsis with an abnormal and unique DSB repair mechanism as the level of RPA2 foci sustained high while those of RAD51 and DMC1 foci dropped significantly compared with the other two types of zygSC-LCs.

### BEND2 interacts with transcriptional suppressors

As BEN proteins were predicted to mediate protein–protein and protein–DNA interactions, and since supportive evidence has been acquired for several family members, we next applied co-IP-mass spectrometry (co-IP-LC-MS/MS) to identify potentially interacting partners of BEND2. Co-IP was conducted using mmAb-FLAG to pull down FLAG-BEND2 and its interacting partners from testicular lysates from FLAG-BEND2 KI mice, and testicular lysates from WT mice were used as negative controls. By analyzing proteins enriched in FLAG-BEND2 KI samples in three independent experiments, we identified several potential BEND2-interacting proteins (Fig. 5A).

**Fig. 5.**
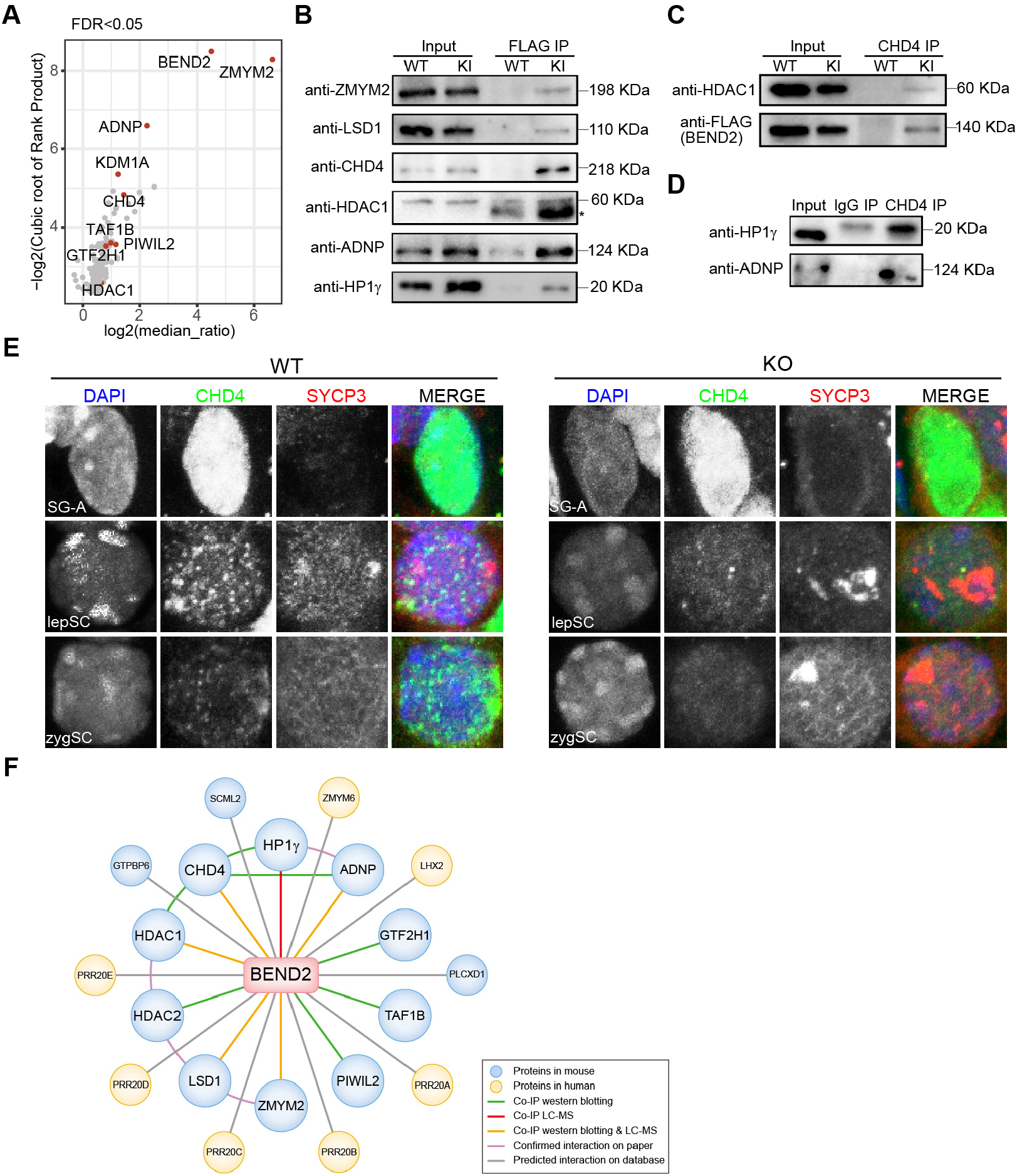
MS analyses of BEND2-interacting factors in testicular extracts. **(A)** LC-MS/MS analysis of enriched protein from co-IP. Protein identification was performed in the presence of wash buffer containing 300 mM or 500 mM NaCl; WT mice served as background controls (n=3 independent biological replicates, with each replicate containing two 15-dpp KI mice). **(B)** Co-immunoprecipitation of BEND2 with ZMYM2, LSD1, CHD4, HDAC1, ADNP, and HP1γ in testis from WT and *Bend2*^FLAG/Y^ mice at 15 dpp. **(C)** Co-IP western blotting analysis used to confirm the interaction of CHD4 and BEND2 or HDAC1 in the testis. **(D)** Co-IP western blotting analysis to confirm the interaction of CHD4 and HP1γ or ADNP in the testis. **(E)** Immunofluorescence of testicular sections with SYCP3 (red) and CHD4 (green) in WT and KO mice, and DNA was counterstained with DAPI (blue); their merged images are shown (scale bar, 10 μm). **(F)** The network of BEND2-interacting proteins.

FLAG-BEND2 manifested the highest enrichment rank among all enriched proteins, indicating that our method was reliable. The next top-three most significantly enriched proteins were ZMYM2, ADNP, and KDM1A (also known as LSD1). ZMYM2 is a member of the MYM (myeloproliferative and mental retardation)-type zinc finger protein family that contains six members in the human and mouse genomes (*31*). LSD1 is the first histone demethylase to be discovered and removes methyl groups from H3K4me or H3K4me2 (*32*). ZMYM2 has been identified as a component of LSD-containing repressive complexes, including the nucleosome remodeling and histone deacetylase (NuRD) complex (*33–35*). These complexes typically also contain histone deacetylases such as HDAC1 and HDAC2 that act upstream of LSD1 (*36*). Intriguingly, we found that HDAC1 and 2 were indeed enriched 1.6- and 1.2-fold, respectively, in our FLAG-BEND2 KI samples (Table S1). ADNP is a transcription factor that contains nine zinc fingers and a homeobox domain, and is essential for embryonic and brain development (*37, 38*). It was reported that ADNP, chromatin remodeler CHD4 (which is the motor component of the NuRD complex), and chromatin architectural proteins HP1β and HP1γ formed a stable complex named ChAHP that represses the expression of lineage-specifying genes in ESCs (*39*). CHD4 consistently ranked No. 7 in the list of BEND2-interacting proteins based upon our results. Other potential interacting proteins that have been uncovered include TAF1B, GTF2H1, and PIWIL2 (*40–42*).

The interactions of BEND2 with ZMYM2, ADNP, LSD1, CHD4, HDAC1, and HP1γ were confirmed by co-IP-western blotting results using the testicular lysates from FLAG-BEND2 KI mice (Fig. 5B). As positive controls, the interactions between CHD4 and HDAC1, HP1γ, or ADNP in the testis were also confirmed by our experiments (Fig. 5C, D). We found that CHD4 was abundantly expressed in the nuclei of SG-A and weakly expressed in the nuclei of lepSCs and zygSCs as granules similar to the pattern for BEND2. In *Bend2* KO mice, the signals for CHD4 in lepSCs and zygSC-LCs were dramatically reduced (Fig. 5E). By mining the human PPI data generated using the yeast two-hybrid technique, we also found that human BEND2 might interact with ZMYM6, LHX2, SCML2, GTPBP6, and PRR20A-E (*43*). Therefore, BEND2 appears to interact with a large number of chromatin-binding proteins that are epigenetic regulators and/or transcription factors (Fig. 5F).

### BEND2 preferentially binds to simple sequence repeats

To characterize BEND2-binding sites on the genome, we conducted ChIP-seq analyses by using testicular lysates from FLAG-BEND2 KI mice; and based on the data from six independent experiments, a total of 16,477 peaks were identified (Fig. 6A, S6A, B; Table S2)—and some peaks were validated by ChIP-PCR. (Fig. S6C). Of note, BEND2 peaks were enriched in proximal promoters (from −1 kb to +100 bp of transcriptional start sites), CpG islands, 5’UTRs, and repetitive sequences (*p*<0.05) (Fig. 6B, S6D). Almost all peaks (95%) were localized to the intergenic regions (58%) and introns (37%), and these peaks were enriched in simple repeats, low-complexity sequences, and satellites (Fig. 6B, S6D). The order of enrichment-fold values (ratios of observed-to-expected peak numbers) for the enriched genomic regions were simple repeats (15.9), low-complexity sequences (11.0), satellites (1.8), 5’UTRs (1.7), and promoters (1.3). We were interested in whether BEND2 peaks were enriched with any known or novel motifs, and noted that several similar GA-rich motifs were enriched in BEND2 peaks (Fig. 6C, S6E). The top enriched motif (AGGAC/T/AAGGAC/T/AAG) was present in 44% of peaks (P = 1 × 10^−5036^) (Fig. 6C left), and the average intensity (566 reads/peak) of peaks containing the top motif was 2.5-fold higher than peaks without the motif (P = 1 × 10^−267^). We further conducted motif-enrichment analyses on peaks that contain the top motif and we identified a similar top motif (AA/CG/CGAAAGGAA/TA), and several other known ones (Fig. 6C right, S6E). We observed that this new top motif was similar to the motif for UME1, a protein that associates with histone deacetylases to repress meiotic gene expression during vegetative growth in yeast (*44, 45*); and to the motif for PU.1, a well-known master regulator and pioneer factor in hematopoiesis from the ETS transcription factor family (*46*)(Fig. S6E).

**Fig. 6.**
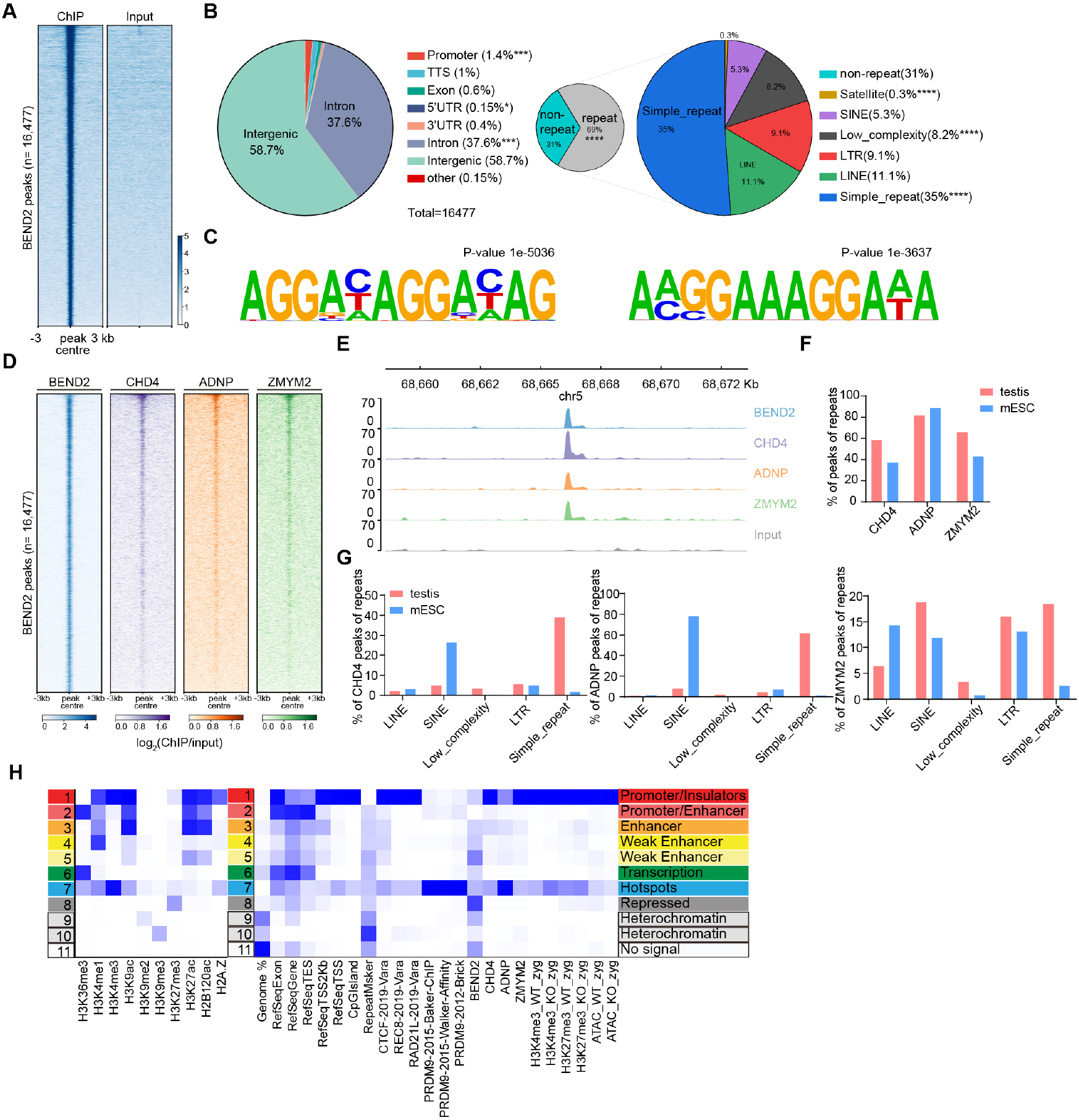
BEND2 binds to multiple chromatin states. **(A)** Heatmap of BEND2 ChIP-seq enrichment across all significant peaks (n=16,477) in the mouse genome. Each row represents a 6-kb window centered on BEND2 peak midpoints, sorted by the BEND2 ChIP signal. Input signals at the same position are shown on the right (average peak intensity of n=6 biological replicates). **(B)** BEND2-binding sites were classified by their genomic locations and repeat types as indicated. **(C)**. The top BEND2 DNA-binding motif predicted by HOMER (left); sequences with the top motif re-analyzed by HOMER (right). **(D)** Heatmap of BEND2, CHD4, ADNP, and ZMYM2 ChIP-seq enrichment across all BEND2 peak midpoints; each raw datum represents a 6-kb window centered on BEND2 peak midpoints. **(E)** Browser view showing ChIP-seq signals of BEND2, CHD4, ADNP, and ZMYM2 co-binding sites. **(F)** Comparison of repeat percentages for CHD4, ADNP, and ZMYM2 in testis and ESC. **(G)** Enrichment comparison of different repeat classes peaks in testis and ESC with respect to CHD4, ADNP, and ZMYM2. **(H)** Heatmap of chromatin states produced by ChromHMM based on 10 histone modifications (left); heatmap showing different enrichment for indicated annotations for each state (right).

We additionally generated ChIP-seq data for the BEND2-interacting proteins CHD4, ADNP, and ZMYM2, and found that reads of these BEND2-interacting proteins were enriched at BEND2 peaks (Fig. 6D, E; Table S3). Similar to the case for BEND2, these proteins preferentially bound to proximal promoters, CpG islands, 5’UTRs, and repetitive sequences that included simple repeats, low-complexity sequences, and satellites (Table S4). Several differences in genomic-region enrichment were observed for BEND2 and its interacting proteins. First, enrichment-fold values vary. For example, BEND2 peaks were only slightly enriched in proximal promoters 1.3-fold while CHD4, ADNP, and ZMYM2 were enriched in these regions 23-, 7-, and 27-fold, respectively. Moreover, BEND2 was enriched in CpG islands two-fold while CHD4, ADNP, and ZMYM2 were enriched 52-, 19-, and 50-fold, respectively. Second, while all of these proteins were only slightly enriched in repeats in general (fold-values no greater than 2), they were enriched in specific repeat-types to greater extents (Fig. 6F, G): fold-values for BEND2, CHD4, ADNP, and ZMYM2 in simple repeats were 16, 17, 28, and 8, respectively. ZMYM2 was also enriched in SINEs and LTRs in addition to simple repeats, low-complexity repeats, and satellites. When we examined the genomic distribution of CHD4, ADNP, and ZMYM2 in mESCs using published datasets (Fig. 6F, G) (*39, 47, 48*), we observed that these proteins were also enriched in promoters, CpG islands, and 5UTRs—except that ADNP was not enriched in 5’UTRs. Moreover, they were enriched in specific repeat types: for example, CHD4 and ADNP were enriched in SINEs and satellites, and ZMYM2 was enriched in SINEs, LTRs, and simple repeats. Notably, almost all BEND2 peaks were enriched in CHD4, ADNP, and ZMYM2 reads, and almost all CHD4 peaks were enriched in ADNP and ZMYM2 peaks, but fewer than half of the CHD4 peaks were enriched in BEND2 reads (Fig. 6D, S6G). Collectively, these data suggested that BEND2 preferentially targets a fraction of its interacting proteins to particular regions (such as simple repeats) in spermatogenic cells.

### BEND2 binds to multiple chromatin states

We were interested in discerning how BEND2 sites in the mouse genome were related to different epigenetic markers such as histone modifications. Spruce et al. (*7*) recently defined an 11-state epigenomic map of leptotene and zygotene spermatocytes by using ChIP-seq data of 10 histone modifications and variants. This was accomplished by using chromHMM, a software package that was based upon a multivariate Hidden Markov Model (*49, 50*). We reproduced the 11-state map and further annotated the map with more genomic features (RefSeq Genes, CpG islands, and repetitive sequences) and the binding sites of proteins involved in meiosis (PRDM9, REC8, RAD21L, and CTCF) (Fig 6H). Consistent with what Spruce et al. found using a single dataset, we found that PRDM9 sites from three independent datasets were mostly enriched in state 7 (recombination hotspots), which is characterized by the moderate-to-high levels of H3K4me3, H3K36me3, H3K4me1, and H3K9ac. REC8, RAD21L, and CTCF were most enriched in state 1, which represents promoters and insulators, and second most enriched in hotspots. We also mapped BEND2 sites to the state map (Fig. 6H). BEND2 was notably enriched in multiple states, with the highest enrichment in state 8; this is typical of H3K27me3, which is a marker for the Polycomb repressive complex-repressed region. These data suggest that BEND2 thus occupies multiple functions in meiosis, one of which might be transcriptional suppression. ChromHMM results also showed that BEND2 and its interacting partners CHD4, ADNP, and ZYMM2 were enriched in state 1, which is annotated as promoters/insulators. These proteins were additionally enriched in state 7, and slightly in states 3–5, which represent enhancers. It is noteworthy that the most enriched states of diverse proteins are usually different, indicating that they do not always remain together. That BEND2 is more or less enriched in many states in spermatocytes suggests that it could be a multifunctional participant in meiosis, and that it likely interacts with different partners in a context/state-dependent manner (Fig. 6H, S6H).

### BEND2 regulates the expression of a large number of genes

We next executed RNA-seq analyses to identify genes that were regulated by BEND2. We identified 2521 and 3918 genes that were up- and downregulated (q < 0.05, n=4), respectively, by *Bend2* KO in adult lepSCs and zygSCs (Fig. 7A, S7A and B; Table S5). With an FDR of no more than 0.05, the upregulated genes were enriched in GO terms related to gene regulation (“transcription,” “mRNA transport,” “RNA processing,” “RNA splicing,” “gene silencing,” “chromatin modification,” “regulation of translation,” “methylation”), DNA activities (“DNA replication,” “DNA repair,” “double-strand break repair,” “nucleosome assembly”), and cell cycle (“cell cycle,” “cell division,” “mitotic nuclear division,” “cell proliferation”) (Fig. 7B, Table S5). Such enrichments are typical of genes that are highly expressed in spermatogonia (*26, 51, 52*), and that are described as “somatic/progenitor program” genes by Hasegawa et al. as they are commonly active in somatic lineages and mitotic phases of spermatogenesis-progenitor cells (*19*). Some of these upregulated genes (*Ehmt2*, *Nanos3*, *Mtor*, *Tdrd1*, *Dazl*, *Lin28a*, *Wdr81*, *Msh2*, *Ercc1*, *Kit*, *Asz1*, *Dnd1*, *Mov10l*, *Bax*, *Piwil2*, *Stra8*, *Kmt2d*, *H3f3b*, *Sohlh1*, *Src*, *Sohlh2*, *Adrm1*, and *Trip13*) are annotated as “germ cell development” or “oogenesis” (Table S5). GO terms enriched in the downregulated genes were quite different, and they were mostly related to meiotic or post-meiotic activities such as “spermatogenesis,” “sperm motility,” “cell differentiation,” “cilium,” and “capacitation.” By replotting the expression of these genes using our previous RNAs-seq data, we found that the upregulated genes were indeed expressed at higher levels in spermatogonia than in spermatocytes, while the downregulated genes exhibited the opposite expression pattern (Fig. S7D). Therefore, it appears that one of the BEND2 functions is to terminate the somatic/progenitor program and to promote the expression of meiotic and post-meiotic genes.

**Fig. 7.**
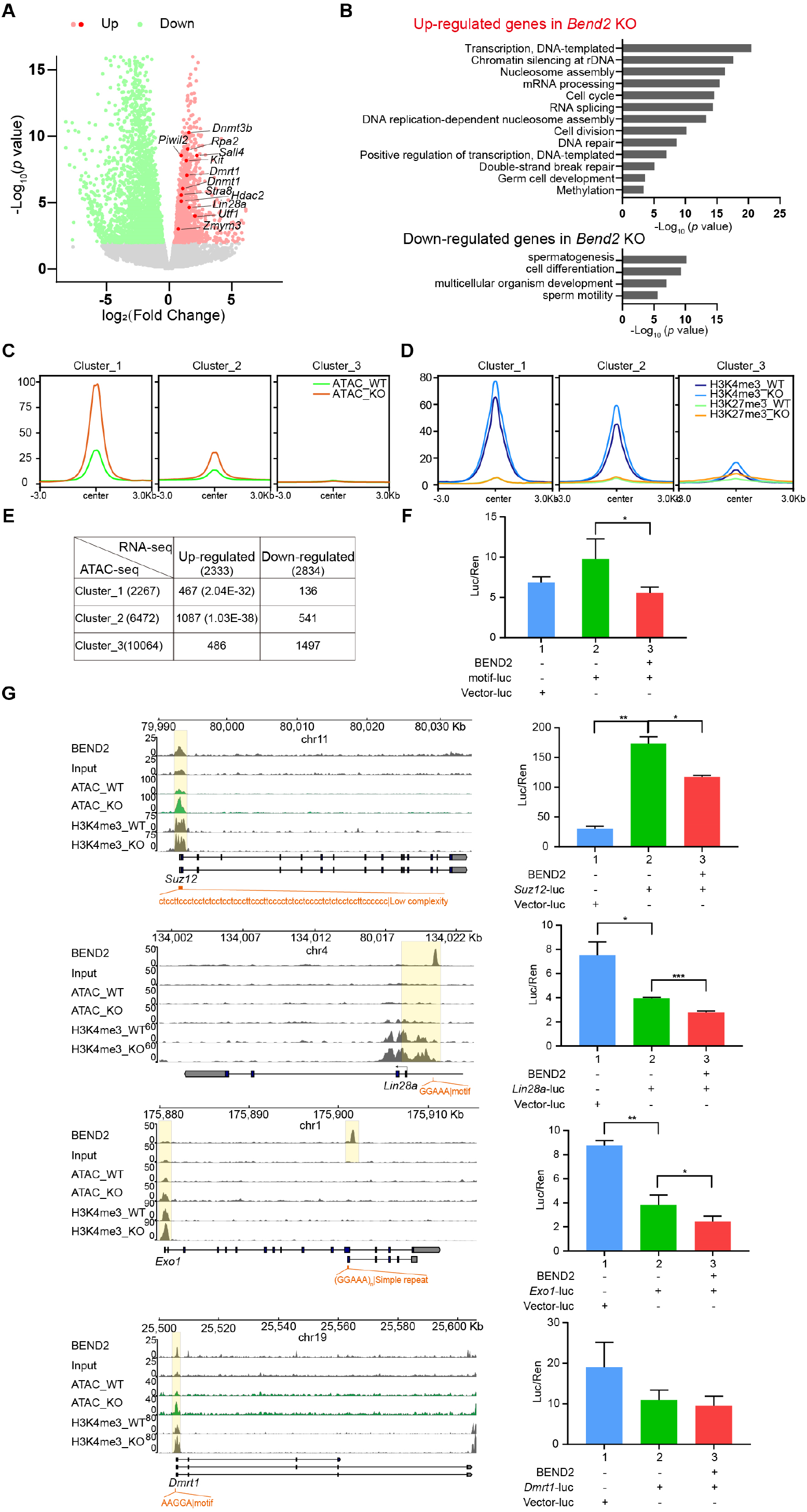
Transcriptomic and epigenetic states change after *Bend2* knockout. **(A)** Volcano plot of transcript levels between cells from adult WT and *Bend2*^-4k/Y^ mice using lepSCs and zygSCs. The differentially expressed genes are highlighted in red (upregulated in *Bend2*^-4k/Y^) and green (downregulated in *Bend2*^-4k/Y^). **(B)** Representative Gene Ontology (GO) terms of the biological process categories enriched in differentially expressed genes. **(C)** Average distribution of ATAC-seq signal around the TSS of three clusters of genes. **(D)** Average distribution of H3K4me3 and H3K27me3 around the TSS of three clusters of genes. **(E)** Correlation analysis between the three clusters of genes by ATAC-seq and differentially expressed genes. **(F)** Validation of DNA-binding motif of BEND2 using dual-luciferase assay. \**p*<0.05 **(G)** Browser view showing BEND2 ChIP-seq, ATAC-seq, and H3K4me3 ChIP-seq signals of target genes (left). Dual-luciferase assay showing the repression of BEND2 target genes (n=3, \*\**p*<0.01 [right]).

We next investigated whether BEND2 regulated gene expression by affecting chromatin accessibility and/or modifications. We isolated zygSCs to implement ATAC-seq and ChIP-seq analyses for H3K4me3 and H3K27me3, and to compare differences in the distributions and intensities of the three signals between WT and *Bend2* KO samples. From the chromHMM state map, we observed that all three types of peaks in both WT and KO mice were mostly enriched in promoters followed by hotspots (states 1 and 7) (Fig. 6H). We did not note any changes in their global enrichment patterns by *Bend2* gene KO from the state map. However, by examining the signal intensities around transcription start sites (TSSs), we observed that ATAC-seq signals were enhanced in a large number of genes (clusters 1 and 2 versus cluster 3) in KO mice (Fig. 7C, Fig. S7H), and the H3K4me3 but not H3K27me3 signals were also enhanced in clusters 1 and 2 (Fig. 7D, Fig. S7I). When we assessed whether up- or downregulated genes based on RNA-seq were enriched in any of these three clusters, we found that only the upregulated genes were enriched in clusters 1 and 2 (P = 2.0 x 10^−32^ and 1.0 x 10^−38^, respectively) (Fig. 7E). Therefore, it appeared that gene repression by BEND2 was achieved by its contribution to maintaining low levels of chromatin accessibility and H3K4me3.

Finally, we carried out luciferase assays to examine whether BEND2 suppressed gene expression by binding to genomic regions identified by the ChIP-seq analyses (Fig 7F-G). We first synthesized a DNA fragment containing five copies of the GGAAA consensus motif and inserted it upstream of the basal promoter of the luciferase-expressing plasmid. Intriguingly, the 5xGGAAA sequence enhanced the promoter activity by itself, and BEND2 enabled this enhancement to revert to basal levels. Based on the ChIP-seq data, we next tested several putative native BEND2-binding sites that were located either around the TSS (*Suz12*, *Lin28a*, *Dmrt1*) or in the gene body (*Exo1*) of genes that were upregulated in *Bend2* KO mice, and we observed that the overlapping or nearby ATAC-seq and H3K4me3 peaks were also upregulated in KO mice. BEND2 significantly reduced the activities of three of the four putative binding sites (*Suz12*, *Lin28a*, *Exo1*), regardless of whether the sites by themselves augmented or attenuated the activity of the basal promoter in the luciferase plasmid (Fig. 7G). These results further supported the concept that BEND2 functions as a transcriptional suppressor of certain genes by binding to and modifying the chromatin accessibility and histone modifications of particular genomic regions.

## DISCUSSION

Meiosis is a highly complex process that entails numerous concurrent or sequential steps that must be coordinated by a large number of regulators. And investigators have in recent years repeatedly described novel regulators of meiosis using phenotypic evaluation of their gene KO mice via the highly efficient CRISPR/Cas9-based gene-editing technology. In the present study, we identified BEND2 as another novel regulator that is specifically expressed in male germ cells shortly before and after meiosis initiation, and that is essential for DSB repair and synapsis using gene KI and KO mice. We also demonstrated that BEND2 interacts with other chromatin-binding/regulating proteins and regulates chromatin state and transcription. Our work has thus contributed another significant component to the arcane and complex physiologic process that is meiosis.

### The BEN protein family

The BEN protein family is a relatively new family that was discovered using bioinformatic analyses, and studies on this family are limited. BANP (BTG3 associated nuclear protein, also known as scaffold/matrix-associated region 1 (SMAR1, or BEND1), has been reported to act as both a tumor suppressor and immunomodulator (*53*), and to repress cyclin D1 expression by recruitment of the SIN3/HDAC1 complex to its promoter and to direct histone modifications from a distance (*54*). E5R is a virosomal protein from the chordopoxvirus subfamily and likely plays a role in organizing viral DNA during replication or transcription. NAC1 (nucleus accumbens-associated protein 1, also called NACC1 or BEND8) participates in various biological processes that include neuronal activity, pluripotency of ESCs, and tumor growth, and it interacts with HDAC3, HDAC4, and REST corepressor 1 (CoREST) (*55, 56*). NAC1 was recently reported to bind DNA directly through the BEN domain in a sequence-specific manner (*57*).

BEND3 contains four BEN domains; is associated with HP1α, HP1β, HP1γ, and H3K9me3-containing heterochromatic foci; and represses transcription through interactions with HDAC1, 2, 3, and SALL4—a transcriptional repressor that also associates with the NuRD complex (*58*). In the absence of DNA methylation or H3K9me3 in mouse ESCs, BEND3 recruits the MBD3/NuRD complex to pericentromeric regions and is necessary for PRC2 recruitment and H3K27me3 establishment at major satellites, suggesting that it is a key factor in mediating a switch from constitutive to facultative heterochromatin (*59*). BEND3 also represses rDNA transcription by interacting with the nucleolar remodeling complex (NoRC) (*60*) and with Suv4–20h2, an enzyme responsible for H4K20 trimethylation (*61*). BEND5 and BEND6 contain a single BEN domain and are therefore (together with three fly proteins) called “BEN-solo” factors (*62*). BEND6, similar to its *Drosophila* homolog Insensitive, is most abundantly expressed in the brain and inhibits Notch target genes (*63, 64*). Importantly, BEND5 and the fly BEN-solo factors bind directly to specific DNA motifs through their BEN domains (*62, 65*). BEND9 (NACC2, RBB) recruits the NuRD complex to the internal promoter of HDM2 and inhibits the expression of HDM2, an E3 ligase that specifically targets p53 for ubiquitination and subsequent degradation (*66*). These studies on the BEN family members have revealed the following: 1) the BEN proteins tend to interact with a variety of proteins, presumably in a context-dependent manner; 2) most of the interacting partners are components of transcription-repressive complexes involved in chromatin remodeling and/or modification; and 3) the BEN proteins can also act as sequence-specific DNA-binding proteins.

We have uncovered little information regarding BEND2, 4, and 7. Several studies showed that in-frame MN1-BEND2, EWSR1-BEND2, and CHD7–BEND2 gene fusions were detectable in brain and pancreatic neuroendocrine tumors (*67–69*). The MN1 (meningioma 1) gene is a proto-oncogene that encodes a transcriptional regulator, and its mutations and abnormal expression are frequently detected in tumors (*70*). EWSR1 (EWS RNA-binding protein 1) is a multifunctional protein that regulates transcription and RNA splicing, and occupies diverse roles in various cellular processes and organ development— including meiosis (*71*). And CHD7 (chromodomain helicase DNA-binding protein 7) is a chromatin-remodeling enzyme involved in differentiation and transcriptional regulation (*72, 73*). The current observations that all fusion partners of BEND2 are transcriptional regulators and that the fusion proteins maintain the BEN domain of BEND2 suggest that BEND2 probably binds DNA through its BEN domain.

### BEND2 protein and its expression

In the present study, we found that BEND2 was specifically expressed in male germ cells after birth. This conclusion was supported by immunostaining results where we used a rabbit polyclonal antibody against a BEND2-specific polypeptide outside of the two BEN domains, and a mouse monoclonal antibody against the 3xFLAG tag fused to the N-end of BEND2 of the mice with FLAG-BEND2 KI. One interesting observation was that the molecular mass of BEND2 as determined by SDS-PAGE was much greater than expected from the amino acid number. This discrepancy may have resulted from the highly disorganized structure and high hydrophobicity of the predicted protein sequence from bioinformatic analyses. However, we cannot rule out the possibility that the protein was post-translationally modified. The punctate signals of BEND2 in lepSCs (Fig. 1G) that did not co-localize with brighter DAPI signals suggested that BEND2 molecules tend to aggregate in regions that are not typical constitutive heterochromatin. We also failed to detect BEND2 in spermatocyte chromosomal spreads and to express the full-length protein in bacteria, suggesting that the BEND2 protein molecules and their putative associations with each other and with other molecules are likely to be labile.

### The phenotypes of *Bend2* KO mice

As expected from the male germ-cell-specific expression of *Bend2*, the KO mice displayed male infertility without any other overt phenotypes; this conclusion was solidly supported by the observation that all three types of mutant male mice (*Bend2*^-19/Y^, *Bend2*^+1/Y^, *Bend2*^-4k/Y^) were infertile. Detailed phenotypic evaluation was carried out with the *Bend2*^-4k/Y^ males at and after F2 (primarily at F5 when the genetic background was purified to 98% C57/BL6 by repeated crosses between *Bend2*^+/-^ females and C57BL/6J males). Similar to the phenotypes manifested by mice with KOs of many other key meiotic regulators, *Bend2* KO mice exhibited arrested spermatogenesis at the zygonema/pachynema transition, with aberrant DSB repair and chromosomal synapsis. The manifestations of several defects are noteworthy, as they imply molecular functions of BEND2 that warrant further investigation. First, synapsis initiation as marked by the appearance of SYCP1 signals was detectable in a significant portion (∼30%) of the KO lepSCs, while it was rarely observed in WT cells. However, the formation of the invasive single-stranded DNAs essential for recombination and synapsis were basically not affected as indicated by the numbers of RPA, RAD51, and DMC1 foci. Second, non-homologous synapses were frequently seen in KO zygSCs. Non-homologous chromosomal association has been reported in mice with KOs of a number of epigenetic regulators such as SUV39H, DNMT3L, and PRDM9, and was likely caused by the disrupted heterochromatin structure (*14, 15, 74*). Third, inter-sister synapses were common in our *Bend2* KO zygSCs, and inter-sister synapses were observed in cohesion-protein KO mice, including KOs of REC8, SMC1β, and STAG3(*75*). REC8 foci were also consistently reduced in *Bend2* KO mice. The phenotypes of *Bend2* KO mice might therefore reflect those of different meiotic regulators, and this implies that BEND2 is a multifunctional protein that is involved in several processes/steps of meiosis.

### The molecular functions of BEND2

The first clue as to the molecular mechanism underlying BEND2 function in meiosis comes from the observation that it interacts with multiple proteins, of which most are transcriptional repressors that often interact with other BEND proteins. The majority of these proteins (HDAC1, HDAC2, CHD4, LSD1, ZMYM2, HP1g, and ADNP) are either core components or interacting proteins of two repressive complexes: the well-known NuRD complex (*76*) and the recently identified ChAHP complex (*39*). This observation together with the fact that our co-IP washing solution contained 500 mM NaCl suggested that the interactions between BEND2 and these complexes were fairly robust. The overlapping expression windows of BEND2 and CHD4, their similar granular expression patterns, and the reduced signal for CHD4 in *Bend2* KO mice also supported an interaction between these two proteins. The interaction between BEND2 and CHD4 and the fact that CHD4 is involved in regulating DSB repair (*77, 78*) facilitate the clarification of why and how meiotic DSB repair was disrupted in *Bend2* KO mice. Other familiar interacting proteins of BEND2 that we identified in the present study included DNA-binding proteins such as TAF1B, GTF2H1, and PIWIL2. Based on our data and bioinformatic predictions from the PPI database, BEND2 interacts with many other proteins; however, the significance of these interactions is currently unclear. Nevertheless, these observations suggested that BEND2, like other family members, may be important in regulating chromatin activities such as heterochromatin formation/maintenance, transcription and higher-order structure.

The potential regulatory roles of BEND2 with respect to chromatin are also supported by the distribution of its ChIP-seq peaks in various genomic regions and the chromatin states as defined epigenetically. The peaks were enriched in regulatory regions such as promoters, CpG islands, enhancers, recombination hotspots, and PRC-repressed sites. It was particularly interesting that the intronic and intergenic peaks (which comprised 96% of the total peaks) were highly enriched in simple repeats and low-complexity repeats, and that a GA-rich motif was also enriched in these peaks. Compared with other chromatin regulators such as CHD4, ADNP, and ZMYM2 that were mainly enriched in promoters and hotspots (Fig. 6H), BEND2 was enriched in almost all of the regulatory regions and heterochromatin types. BEND2 was only slightly enriched in proximal promoters (1.3-fold), while the other factors were highly enriched (23-, 7-, and 27-fold for CHD4, ADNP, and ZMYM2, respectively). In contrast, enrichment of these proteins in simple repeats was comparable (16, 17, 28, and 8, respectively). However, the types of simple repeats in which CHD4, ADNP, and ZMYM2 were enriched in spermatocytes were somewhat different from those in mESCs. These observations suggested that BEND2, as a germ cell-specific chromatin-binding protein, either guides or stabilizes the genomic distribution of its interacting partners. More evidence for this hypothesis will only be acquired from future studies by examining the distributional changes in these partners in *Bend2* KO cells or in mESCs in which exogenous BEND2 was expressed.

### Regulation of gene expression by BEND2

The analyses of transcriptomic changes in spermatocytes of *Bend2* KO mice provided further clues regarding the molecular functions of BEND2 as a chromatin regulator. It appears that BEND2—similar to Sex comb on midleg-like 2 (SCML2)— also contributes to shutting down the mitotic program and to activating or enhancing the meiotic and post-meiotic program of spermatogenic cells (*19*). As SCML2 was expressed in cells ranging from undifferentiated spermatogonia to spermatocytes, and as global gene-expression changes were not observed until the spermatocyte stage upon Scml2 KO, these authors proposed that the suppressive process was initiated prior to when the effect became obvious after meiotic initiation. This indicated that some components of the SCML2-involved suppressive machinery were not ready until the late stage of spermatogonial differentiation, and therefore BEND2 may be one of the missing components as its expression was noted just prior to meiotic initiation. Therefore, it would be compelling to investigate the relationship between SCML2- and BEND2-mediated suppressive machineries in the future.

Increasing evidence signifies that gene repression is an essential means of gene regulation in diverse cellular developmental processes. As far as germ cells are concerned, somatic genes are repressed in early-stage primordial germ cells in both sexes (*79*), while the meiotic program is prevented in male gonocytes and spermatogonia by the RA-metabolizing pathway (*80*) and proteins such as NANOS2, DMRT1, and SCML2 (*19, 81, 82*). Thus, the identification of BEND2 as a novel repressive regulator indicates that this regulatory scheme may be much more extensive and complex than previously thought. Since a repressor can specifically activate the expression of genes as an indirect result of the repression of other repressors that target the activated genes, it is not surprising that a large group of genes involved in meiotic and post-meiotic activities of spermatogenic cells can be downregulated upon *Bend2* KO. This suggests that BEND2 contributes to the expression of these genes under normal conditions, and among the genes normally repressed by BEND2 (upregulated genes in *Bend2* KO mice), 109 were negative regulators of transcription according to their GO annotations (FRD=0.04; Table S5, line 37). Repressive regulators that are familiar to us in this list include EHMT1/GLP1, EHMT2/G9A, SALL1, SALL4, DNMT1, DNMT3B, SUV39H2, HDAC2, BEND3, and DMRT1; and some of these are known to be of critical significance to spermatogonial proliferation, differentiation, and meiotic initiation. For example, DMRT1 is a repressor of meiotic initiation as its gene KO in mice initiates meiosis precociously (*82*). PRC2, of which SUZ12 is a core subunit, is required for spermatogonial stem cell maintenance and meiotic progression via repression of somatic and meiotic gene expression (*83*). Moreover, we consistently showed with luciferase assays that the TSSs of both *Dmrt1* and *Suz12* contained BEND2-binding sites that were repressive in the presence of BEND2.

In summary, we identified BEND2 as a germ-cell-specific regulator of meiosis with detailed examinations of its expression and function by using gene KI and KO mice. We also demonstrated the molecular mechanisms underlying BEND2’s action as a chromatin modulator and transcriptional repressor by identifying and characterizing its interacting partners, genomic binding sites, and regulated genes. However, there are many more issues that await clarification in the future. For example, there are the questions of whether BEND2 is critical to female meiosis (oogenesis), whether it acts as a direct modulator of the meiotic machinery independent of its role in transcriptional repression, and whether molecular defects can be detected earlier, prior to meiotic initiation. We posit that the results of our comprehensive present study will establish a solid foundation for such future investigations.

### Materials and Methods Animal care

All animal procedures were approved by the Animal Ethics Committee of the Institute of Zoology, Chinese Academy of Science. The mice were housed in a specific pathogen-free facility with a 12 h:12 h light-dark artificial-lighting cycle, with lights off at 19:00, and were housed in cages at a temperature of 22–24°C. All experiments with mice were conducted in accordance with the Guide for the Care and Use of Laboratory Animal Guidelines.

### RT-PCR

Approximately 50 milligrams of mouse tissue was incubated with 1 ml of TRIzol reagent (Invitrogen Cat. no. 15596-026) and homogenized with a Dounce homogenizer. All liquid was transferred to RNase-free Eppendorf tubes and incubated at room temperature for 5 min. We then added 0.2 ml of chloroform, capped the tubes securely, shook them by hand for 15 s, incubated the tubes for ∼3 min at room temperature, and centrifuged them at 12,000 x g for 10 min at 4°C. Supernatants were transferred to a new tube and RNA was extracted using chloroform followed by isopropanol precipitation. The RNA was dissolved in nuclease-free water (P1195, Promega), and the RNA quality was measured with a Nanodrop 2000. To prepare the cDNA library, total RNAs were reverse-transcribed using a High-Capacity cDNA Reverse Transcription Kit (4368814, Applied Biosystems). The primers we used to detect gene expression levels are listed in Table S7. The following conditions were used for PCR: 94°C, 2 min; 30 cycles of 94°C, 1 min; 60°C, 1 min; 72°C, 40 s; and a final extension at 72°C, 10 min. PCR products were separated on 1.5% agarose gels.

### *Bend2* cDNA clone

Total RNA was extracted from adult mouse testis and reverse-transcription was performed as described above. We used nested PCR for the *Bend2* cDNA clone (two pairs of primers were designed and their sequences are provided in Table S7). The PCR products were purified with agarose gel electrophoresis and recovered with an EasyPure quick gel extraction Kit (M2073, TransGen Biotech). The cDNA was cloned into a pGM-T plasmid (VT202-02, TIANGEN Biotech), and the cDNA sequence was identified by Sanger sequencing.

We identified two transcripts when we attempted to clone the cDNA from mouse testes: one (V1) contained the 14 predicted coding exons, while the other (V2) was missing the 4th exon that corresponded to a 35-aa (105 bp) in-frame deletion in the protein (Fig. S1C). Based on the intensities of the cDNA bands in the stained agarose gel, V1 was more abundant than V2.

### Generation and genotyping of *Bend2* mutant mice

*Bend2* mutant mice were generated through the CRISPR/Cas9 gene-editing approach (*27*). One male and three female founder mice with DBA/2/C57BL/6J background were acquired by using different gRNAs. For *Bend2*^-4k/Y^ mice, we designed two gRNAs for the long-fragment deletion using methods described previously (*84*), while only one gRNA was designed for the generation of *Bend2*^+1/Y^ and *Bend2*^-19/Y^ mice. Genomic DNA extraction followed the standard proteinase-K-chloroform method. Genotyping for *Bend2*^+1/Y^ and *Bend2*^-19/Y^ mice was executed by Sanger sequencing following PCR amplification, while two pairs of primers were designed for genomic identification of *Bend2*^-4k/Y^ mice; p1 and p2 primers were used to identify the WT and KO allele, respectively (all primers are listed in Table S7).

### Generation of anti-BEND2 antibody

Antibodies to mouse BEND2 were produced by ABclonal Technology (Wuhan, China), and generated by immunization of rabbits with the following peptides: 1-30aa (MESDTDDSHISYDGDELFSEDFGSDIEDTS-C), and 585-614aa (DVRESVKRERVDFEHTPDANPEGSDNASIN-C). Antibodies were purified using antigen-specific affinity columns.

### Generation and genotyping of *Bend2-3xFLAG* knock-in mice

*Bend2-3xFLAG* knock-in mice were generated through the CRISPR/Cas9 gene-editing approach (*85*). The targeting fragment was designed to insert 3xFLAG in-frame with the coding sequence just after the first ATG of the *Bend2* genomic locus. To ensure accuracy, we designed two gRNAs and compared their efficiencies: KI-gRNA-2 was more efficient and is listed in Table S7. The donor DNA contained a 3xFLAG-Linker, with left and right homology arms (800 bp). Donor DNA was synthesized by Sangon Biotech. Genomic DNA extraction also following the standard proteinase K-phenol/chloroform method. PCR was then performed to identify genotype (primers are listed in Table S7).

### Histology, hematoxylin and eosin (H&E) staining, immunohistochemistry, immunofluorescence, and TUNEL staining in testicular sections

Testes or epididymides from WT and KO mice were dissected and fixed with Bouin’s solution or 4% paraformaldehyde (PFA), and then embedded in paraffin and sectioned at 5 μm for staining. For H&E staining, Bouin’s solution-fixed sections were stained with H&E following standard protocol. For immunofluorescence or immunohistochemical study, 4% PFA-fixed sections were dewaxed and rehydrated, and then slides were incubated with sodium citrate buffer (pH 6.0) at 95°C for 10 min to retrieve antigen. Five percent BSA or 5% skimmed milk was used to block nonspecific antigens for 1 h at room temperature (RT). Primary antibodies were diluted with 5% BSA and then incubated with sections at 4°C overnight. After washing three times with PBS, diluted secondary antibodies conjugated with fluorescent tag or HRP were incubated with sections. For immunofluorescence, DNA was stained with DAPI diluted by PBS, and photomicrographs were taken with confocal fluorescence microscopes (LSM780, Zeiss; LSM880, Zeiss; Nikon A1 N-SIM S, Nikon). For immunohistochemistry, a DAB solution as chromogen was diluted and used to cover the sections at RT for 10 min; this was immediately followed by PBS to stop the reaction. Slides were dehydrated and nuclei were stained with hematoxylin. Images were taken with an optical microscope (ECLIPSE 80i, Nikon). We implemented the DeadEnd™ Fluorometric TUNEL System (G3250, Promega) for TUNEL staining.

### Preparation of tissue extracts and western immunoblotting analysis

Tissues were harvested and washed once with PBS. After mechanically shearing them into pieces, we transferred the tissues to a Dounce for homogenization in RIPA lysis buffer (P0013B, Beyotime) containing protease-inhibitor cocktail, and the mixture was incubated on ice for 30 min. Cell debris was removed by centrifugation at 13,500 x g for 15 min at 4°C, and lysates were boiled with 5x SDS loading buffer for 10 min. Tissue extracts were electrophoresed on SDS-PAGE gels containing different concentrations of separation gels between 6% and 15% based upon the molecular weights of the proteins, and then blotted onto PVDF membranes (88518, Thermo). Membranes were blocked with 5% skimmed milk for 1 h at room temperature, and then incubated in dilutions of primary antibodies overnight at 4°C. After washing three times with PBST, membranes were incubated in horseradish peroxidase (HRP)-conjugated secondary antibodies (diluted in PBS) for 1 h at room temperature, and the membranes were then washed three times with PBST at room temperature with gentle shaking. The protein blots were ultimately detected with SuperSignal^TM^ West Pico Plus Chemiluminescent Substrate (34577, Thermo) and imaged on a Bio-Rad Universal Hood II imaging system.

### Spermatocyte chromosome spreads and immunofluorescence of spermatocytes

Spermatocyte chromosome spreads of the testicular samples were performed using the drying-down technique (*86*). Briefly, the testes were dissected from 2–4-month-old mice, and the seminiferous tubules were washed in PBS. The tubules were then placed in a hypotonic extraction buffer for 30–60 min. Subsequently, the tubules were minced in 0.1 M sucrose (pH 8.2) on a clean glass slide and pipetted repeatedly to create a cellular suspension; the suspensions were then spread on slides containing 1% PFA and 0.15% Triton X-100 (pH 9.2), and dried for at least two hours in a closed box with high humidity. Finally, the slides were washed twice with 0.4% Photo-Flo 200 (Kodak), dried at room temperature, and stored at −80°C for immunofluorescent staining. Slides were equilibrated to RT, and then each was washed with PBS twice for 5 min with gentle shaking. BSA (5%) was dropped onto the slides for blocking, and they were covered by parafilm for one hour in a humidified box. Fluorescence staining was identical to that described for immunofluorescence staining. Immunolabeled nuclei with chromosomal spreads were imaged on confocal laser scanning microscopes (LSM780, Zeiss; LSM880, Zeiss) using a 63× oil-immersion objective. For SIM, images were taken on a Nikon A1 N-SIM S microscope.

### Co-IP-mass spectrometric analyses

Four testes from 15-dpp *Bend2-3xFLAG* knock-in mice or WT mice were homogenized by using Dounce homogenizers in 1 ml of cold lysis buffer (20 mM Tris-HCl [pH 7.4], 150 mM NaCl, 1 mM EDTA, 5% glycerol, and 1% NP-40, with fresh 100x proteinase inhibitor added just before use), and then incubated on ice for 30 min. We removed cellular debris by centrifugation at 13,500 g for 15 min at 4°C, and cell lysates were precleared with 25 μl of protein G beads (10003D, Invitrogen) at 4°C for one hour. For mass spectrometry, 80 μl of Anti-FLAG Magarose Beads (SM00905, SMART LIFESCIENCES) were added to the precleared lysates and the mixture rotated at 4°C overnight. The beads were washed four times with cold, low-salt co-IP wash buffer (20 mM Tris-HCl [pH 7.4], 300 mM NaCl, 1 mM EDTA, 5% glycerol, and 1% NP-40, with fresh 100x proteinase inhibitor added just before use) or high-salt co-IP wash buffer (20 mM Tris-HCl [pH 7.4], 500 mM NaCl, 1 mM EDTA, 5% glycerol, and 1% NP-40, with fresh 100x proteinase inhibitor added just before use), rotating each time for 15 min at 4°C. Proteins were eluted from the beads with 25 μl of 1x SDS loading buffer and boiled for 10 min. The presence of proteins in the immunoprecipitated samples was confirmed by SDS-PAGE using a 10% concentration of separation gel and sliver-staining. Whole samples collected from the gel were used to perform mass spectrometric analyses. For co-IP western blotting, 80 μl of precleared lysates mixed with 5x SDS loading buffer were boiled for 10 min as an input sample prior to immunoprecipitation. The other lysates were incubated with 5 μg of antibodies or isotype IgG as experimental samples and negative control, respectively, and rotated at 4°C overnight. We then added 30 μl of Dynabeads Protein A/G (10001D, 10003D, Invitrogen) to each sample based on the host species of antibodies, and incubated the samples at 4°C for four hours. The washing and elution steps were the same as described above.

### In-gel digestion of proteins

The protein bands in each lane were cut into small plugs, washed twice with 200 μl of distilled water, dehydrated with acetonitrile for 10 min, and dried in a Speedvac for approximately 15 min. Gel plugs were treated with 10 mM DTT in 25 mM NH_4_HCO_3_ for 45 min at 56°C for subsequent reactions, and alkylated with 40 mM iodoacetamide in 25 mM NH_4_HCO_3_ for 45 min at room temperature in the dark, followed by two washes with 50% acetonitrile in 25 mM NH_4_HCO_3_. The gel plugs were ultimately dried and digested with trypsin (40 ng for each band) in 25 mM NH_4_HCO_3_ overnight at 37°C. We added formic acid to the reaction buffer for a final concentration of 1% in order to stop the enzymatic reaction. The solution was then transferred to a sample vial for LC-MS/MS analysis.

### Rank product analysis and false-positive rate

We determined significance levels for each protein after MS using rank product (RP) analysis (*87*) and false-positive rate (FDR). First, we merged three replicates of MS_ratio values (from our MS results) by replacing missing values with 1 and kept those genes that reflected an MS_ratio in all thrice-repeated MS experiments. We then ranked MS_ratio values from large to small and calculated rank products for each gene using the following formula: Rank Product = (rank1/n) * (rank2/n) * (rank3/n), where n was the total number of genes, rank1 the rank in the first MS experiment, rank2 the rank in the second MS experiment, and rank3 the rank in the third MS experiment. As a result, each gene had a rank product value based on thrice-repeated MS experiments, and as with a significant p-value, the smaller the rank product score, the more significant the gene. Finally, we computed a false-positive rate (FDR) for each rank product with the Benjamini–Hochberg procedure (*88*) using the p.adjust() function in the R program. The input value was the rank product value for each gene, and by setting the parameter “method” as “fdr” and “n” as the total number of genes that we considered, we achieved a false-positive rate for each rank-product value.

### Sample preparation for RNA-seq

We collected leptotene or zygotene spermatocytes via FACS (*89*). Briefly, the testes from one adult WT mouse or from three adult KO mice were digested by two-step methods. Seminiferous tubules were segregated with collagenase I and DNase I, and then 0.25% trypsin and DNase I were used to obtain a single-cell suspension. Testicular cells isolated from WT and KO mice were then sorted by FACS after Hoechst 33342 staining. Different types of spermatocytes were then collected through Hoechst Blue and Hoechst Red channels, and RNA was prepared following the TRIzol (Invitrogen) protocol. All RNA libraries were constructed at the same time using the NEBNext Ultra Directional RNA Library Prep Kit for Illumina (E7760) according to the manufacturer’s recommendations, and oligo (dT) beads (NEB) were used to isolate poly (A) mRNAs.

### ChIP-seq

Approximately 60 mg of testicular tissues from mice at 15 or 40 dpp was disaggregated by Dounce homogenization and the chromatin was crosslinked in PBS containing 1% formaldehyde for 10 min at RT. Fixation of chromatin was halted with a 1.25 M glycine solution and washed three times with cold PBS. Cells was lysed for 30 min on ice by adding cell-lysis buffer (50 mM Tris-HCl [pH 8.0], 10 mM EDTA, and 1% SDS, and fresh 1 mM PMSF and protease inhibitor were added before use). Chromatin was then sonicated for 20 s at 25% power in 30-s pulses for 20 cycles, cellular debris was removed by centrifugation, and the supernatant was precleared by IgG for 2 h at 4°C. About 1/20^th^ of the chromatin was saved as an input sample, and the remainder was diluted to 10x volume with IP dilution buffer (20 mM Tris-HCl [pH 8.0], 150 mM NaCl, 2 mM EDTA, 1% Triton X-100, and 0.01% SDS), and incubated with 10 μg of antibody for 2 h at 4°C. Protein A/G (10001D, 10003D, Invitrogen) Dynabeads were added to capture targeted chromatin overnight. Beads were washed with a four-step wash buffer (low-salt wash buffer, 20 mM Tris-HCl [pH 8.0], 2 mM EDTA, 50 mM NaCl, 1% Triton X-100, and 0.1% SDS; high-salt wash buffer, 20 mM Tris-HCl [pH 8.0], 2 mM EDTA, 500 mM NaCl, 1% Triton X-100, and 0.01% SDS; LiCl wash buffer, 10 mM Tris-HCl [pH 8.0], 1 mM EDTA, 0.25 M LiCl, 1% NP-40, and 1% deoxycholic acid; TET buffer, 10 mM Tris-HCl [pH 8.0], 1 mM EDTA, and 0.1% Tween 20); beads were washed twice for each step. After the chromatin was eluted from the beads by elution buffer (10 mM Tris-HCl [pH 8.0], 1 mM EDTA, and 1% Tween 20), cross-linking was reversed with 5 M NaCl at 65°C for 16 h; and RNA and protein were digested by adding RNase A and protease K for 2 h at 37°C and 45°C, respectively. Extracted DNA was used to construct a library with the NEBNext Ultra II DNA library Prep Kit for Illumina (E7645, NEB), and qualified libraries were sequenced with an Illumina Novaseq 6000 to obtain paired-end 150-nt reads.

### CUT&RUN

We collected zygotene spermatocytes via FACS as described above, and CUT&RUN was primarily performed according to Henikoff et al. (*90*). In brief, before experimentation, 10 μl of concanavalin A beads (BP531, Bangs Laboratories) were washed twice with binding buffer, resuspended with 10 μl of binding buffer, and maintained on ice. For each sample, ∼20,000 cells were suspended in wash buffer, incubated with prepared beads, and mixed for 10 min at RT. After discarding liquids, beads were incubated with antibody buffer on a Thermomixer at 4°C overnight. Beads were washed once with Dig-Wash buffer, resuspended with the same buffer containing pAG-MNase at a final concentration of 700 ng/ml (we purified the pAG-MNase according to methods described previously (*91*), and this mixture was incubated for three hours at 4°C. Beads were then washed twice with the Dig-Wash buffer, resuspended in the same buffer containing 2 mM CaCl_2_, vortexed, and placed on ice as soon as possible. After 30 min, a 2x stop buffer was added to quench the digested reaction, and it was incubated at 37°C for 30 min. The suspensions were collected and the DNA was extracted. We prepared the library using a NEBNext Ultra II DNA library Prep Kit for Illumina (E7645, NEB), and sequenced the qualified libraries with an Illumina Novaseq 6000 system to obtain paired-end 150-nt reads.

### ATAC-seq

Zygotene spermatocytes were collected by FACS as described in CUT&RUN, and cells were washed twice with cold PBS. To prepare nuclei, cells were lysed with cold lysis buffer (10 mM Tris-HCl [pH 7.4], 10 mM NaCl, 3 mM MgCl_2_, and 0.1 % NP-40), maintained on ice for 10 min, and centrifuged at 500 x g for 5 min at 4°C. After carefully removing the suspension, we resuspended the pellet in the transposase reaction mix (TD501, Vazyme) and incubated it at 37°C for 10 min. Fragments were then immediately purified with 2 x AMPure XP beads (A63881, Beckman), and library amplification was performed using the TruePrep DNA Library Prep Kit V2 for Illumina (TD501, Vazyme).

### ChromHMM analyses

Chromatin-state discovery and genome annotation with ChromHMM was carried out by following the protocol by Ernst and Kellis (*50*) using ChromHMM software v1.22. Based on the study by Spruce et al. (*7*) and personal communications with the corresponding author Dr. Christopher L. Baker, we compiled the cellmarkfiletable shown in Table S6. Datasets indicated in the Table were downloaded from GEO and mapped to the mouse genome (mm10) using Bowtie2, and Bam files from sample replicates were merged and binarized. The initial state map we created was compared with the published version, and the correspondences between states in our initial map and the one published (Table 1A by Spruce et al. (*7*)) were established visually; the states in our initial map were then re-ordered to generate the final map by using the “java -mx4000M -jar ChromHMM.jar Reorder” command. Data for other markers that were either published or produced in the present study were aligned to the map by using the “java -mx4000M -jar ChromHMM.jar OverlapEnrichment” command.

### RNA-seq analysis

Our RNA-seq analyses followed a standard procedure that included mapping sequence reads to the mouse genome mm10 by using Bowtie2 and identifying differentially expressed genes by using the DESeq2 R package.

### ChIP-seq and CUT&RUN analysis

ChIP-seq raw reads were trimmed to remove the adapter sequence when converting to a fastq file, and the trimmed ChIP-seq reads were mapped to the UCSC mm10 genome using Bowtie2 (v2.4.1) (*92*) with default parameters. For BEND2, peak calling was performed using Pepr (*93*) with default parameters and the corresponding inputs as background, and peaks that mapped to blacklist (*94*) regions were removed. For ADNP, CHD4, and ZMYM2, peak calling was performed using MACS2 (v2.2.7.1) (*95*) (https://github.com/macs3-project/MACS) with default parameters, except that the *q* value was less than 0.01. Annotation of genomic locations and repeat types were generated using HOMER (v4.11), and heatmaps were generated using the command-line version of deepTools (v3.5.0) (*96*). Distribution of BEND2-binding sites on the chromosomes were generated by ChIPseeker (v1.24.0) (*97*), and HOMER (v4.11) was used with default settings to identify enriched motifs in BEND2 peaks. RepeatMasker and TSS-location files were downloaded from the UCSC website, and we achieved a Genome Browser view of the NGS data by using the command-line version of pyGenomeTracks.

### ATAC-seq analysis

Paired-end reads were aligned with Bowtie2 using default parameters, and only uniquely mapping reads were retained for further analysis; PCR duplicates and blacklist-region reads were removed. Peak calling was executed using MACS2 (v2.2.7.1). Different gene clusters and heatmaps were then generated using the command-line version of deepTools (v3.5.0), and correlation analysis between the three clusters of genes by ATAC-seq and differentially expressed genes was based upon a hypergeometric distribution.

### Luciferase assay

*Bend2* cDNA was cloned into a pFLAG-CMV-4 vector, and the BEND2-targeted regions for *Dmrt1*, *Suz12*, *Lin28a*, and *Exo1* were PCR-amplified from mouse genomic DNAs isolated from mouse tail tips and cloned into a PGL4.23-luciferase vector (Promega, E8411). Five copies of a GGAAA sequence were synthesized by Sangon Biotech and cloned into a PGL4.23-luciferase vector. TF-expressing plasmids, promoter-luciferase plasmids, and the pRL-TK-*Renilla* constructs as internal controls were co-transfected into 293FT cells on 96-well plates using X-Transcell reagent (bjyf-Bio technology) following the manufacturer’s protocol. Cell extracts were prepared 48 h after transfection using the lysis buffer provided in the Dual-Luciferase Reporter Assay System kit (Promega), and luciferase activity was measured on a Synergy Neo2 Multi-Mode Microplate Reader instrument (Bio-Tek) according to the manufacturer’s protocol. *Renilla* luciferase activity was used to normalize the firefly luciferase activity.

### Statistical analysis

All experiments reported herein were independently repeated at least three times, and all values in the Figures are depicted as mean ± SEM unless stated otherwise. We used Excel 2016 or GraphPad Prism 7 to perform statistical analyses. To analyze the differences between two groups, we used two-tailed unpaired Student’s *t*-tests. To examine whether a group of genes (or genomic features) classified upon one parameter were enriched with a group of genes classified upon another parameter, we executed the R function phyper (k-1, M, N-M, n, lower.tail =FALSE), where N was the total number of genes, M was the number of genes that were positive for the second parameter, n was the number of genes that were positive for the first parameter, and k was the number of genes that were positive for both parameters. This function was based on a hypergeometric distribution. For statistical analysis of focus number comparisons of RPA2, DMC1, and RAD51, data were analyzed with the Tukey multiple-comparison test after one-way ANOVA. For statistical analysis of inter-REC8 distance comparisons, *p* value was obtained with two-tailed, unpaired t-test. No samples or animals were excluded from analyses, sample-size estimates were not used, and the mice analyzed were litter mates. Investigators were not blinded to mouse genotypes or cell genotypes during experiments. For all figures, *, **, and *** represent *p* < 0.05, *p* < 0.01, and *p* < 0.001, respectively. NS (not significant) indicates not statistically significant (i.e., *p* > 0.05).

## Acknowledgments

We thank Mengcheng Luo (Wuhan University) for antibodies and critical suggestions. We thank Jifeng Wang and Xiang Ding in Institute of Biophysics, Chinese Academy of sciences for their technical assistance. We thank Shiwen Li, Xili Zhu, Xia Yang, and Qing Meng in Institute of Zoology, Chinese Academy of Sciences for their technical assistance. We thank LetPub (www.letpub.com) for its linguistic assistance during the preparation of this manuscript.

## Founding

This work was supported by the Ministry of Science and Technology of China (2018YFE0201100 to C.H. and 2016YFC1000606 to C.H.) and the National Natural Science Foundation of China (31771631 to C.H. and 31970795 to C.H.).

## Author contributions

Conceptualization: C.-S.H, S.-G.D, L.-F.M, D.X. Methodology and Investigation: L.-F.M, D.X, H.-Y.N, J.C. Visualization: L.-F.M, X.-W.L, D.X, C.-X.G. Supervision: S.-G.D and C.-S.H. Writing—original draft: L.-F.M, D.X, C.-S.H. Writing—review & editing: C.-S.H.

## Data and materials availability

All data needed to evaluate the conclusions in the paper are present in the paper and/or the Supplementary Materials. Additional data related to this paper may be requested from the authors.

