## Supplemental figure for "Identification and characterization of BEND2 as a novel and key regulator of meiosis during mouse spermatogenesis"

**This PDF file includes:**

Fig. S1 Identification of the germ cell-specific factor BEND2.

Fig. S2 *Bend2* is essential for spermatogenesis.

Fig. S3 Persistence of γH2AX in *Bend2* knockout spermatocytes.

Fig. S4 BEND2 is essential for chromosomal synapsis in meiosis.

Fig. S5 The immunofluorescence pattern of CHD4 is altered in *Bend2*$^{-4k/Y}$ mice.

Fig. S6 Characteristic analysis of the BEND2-binding site.

Fig. S7 Transcriptome analysis of spermatocytes in meiotic prophase I from *Bend2* KO mice.

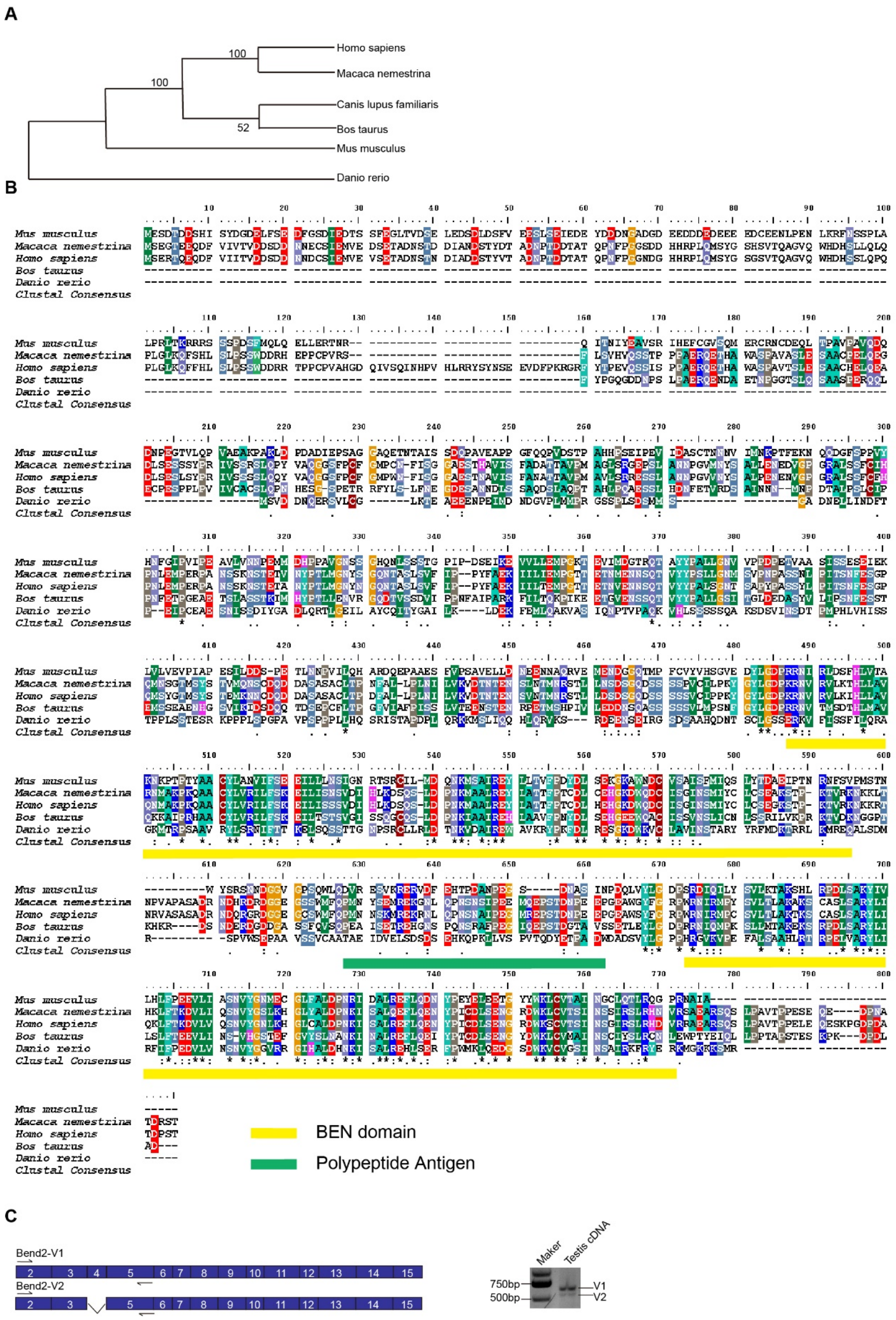
A
Homo sapiens
Macaca nemestrina
Canis lupus familiaris
Bos taurus
Mus musculus
Danio rerio
100
100
52
B
Mus musculus
Macaca nemestrina
Homo sapiens
Bos taurus
Danio rerio
Clustal Consensus
BEN domain
Polypeptide Antigen
C
Bend2-V1
Bend2-V2
Maker
Testis cDNA
750bp
500bp
V1
V2

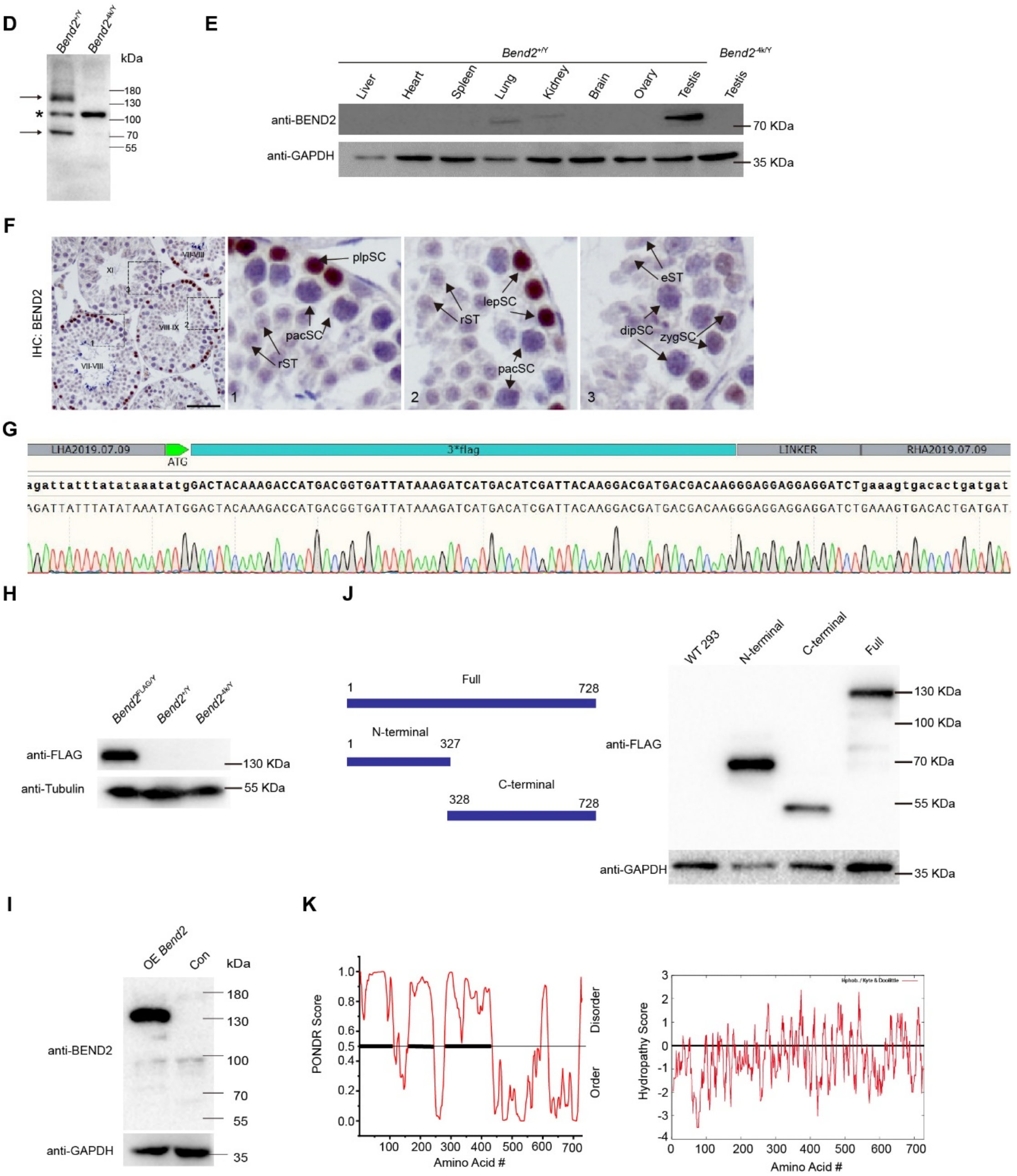


**Fig. S1 Identification of the germ cell-specific factor BEND2.**

**(A)** The phylogenetic tree for BEND2 protein among various vertebrates. **(B)**. Amino acid-sequence alignment of BEND2 protein among various vertebrates and using the ClustalW2 multiple-alignment program revealed high conservation among vertebrates in the BEN domain (" *",":" and "." represent the degree of conservation, and "*" denotes the highest conservation). The green region is the polypeptide selected for the preparation of polyclonal antibodies and the yellow regions represent the two BEN domains. **(C)** Two transcripts of *the Bend2* gene in mouse testes were detected during cDNA cloning; the left plot indicates the structure of exons in the two transcripts, and the right figure shows the expression levels of the two transcripts. **(D)** Western blotting indicated two BEND2-related proteins in mouse testicular lysates using rpAb-B2. **(E)** Western blotting analyses of BEND2 expression in multiple mouse organs using rpAb-B2. **(F)** Immunohistochemical staining of BEND2 in testicular sections of various seminiferous stages using rpAb-B2. **(G)** Sanger sequencing of the *Bend2*$^{FLAG/Y}$ mouse genome. **(H)** Western blotting detection of FLAG-BEND2 expression among *Bend2*$^{FLAG/Y}$, *Bend2*$^{+/Y}$, and *Bend2*$^{-4k/Y}$ testicular lysates using mmAb-FLAG. **(I)** Western blotting detection of FLAG-BEND2 expression in lysates from FLAG-BEND2- overexpressing 293FT cells using rpAb-B2. **(J)** Western blotting analyses of full length, N-terminal, and C-terminal proteins in lysates of plasmid-transfected 293FT cells using mmAb-FLAG. **(K)** Characteristic analysis of the BEND2 protein sequence; the left diagram shows disordered analysis of BEND2; the black line is a predictor of the natural disordered regions (PONDR) score for intrinsic disorder, with >0.5 considered to be disordered; the right diagram indicates hydropathic analysis of BEND2, and the red line depicts the hydropathy score (positive is hydrophobic).

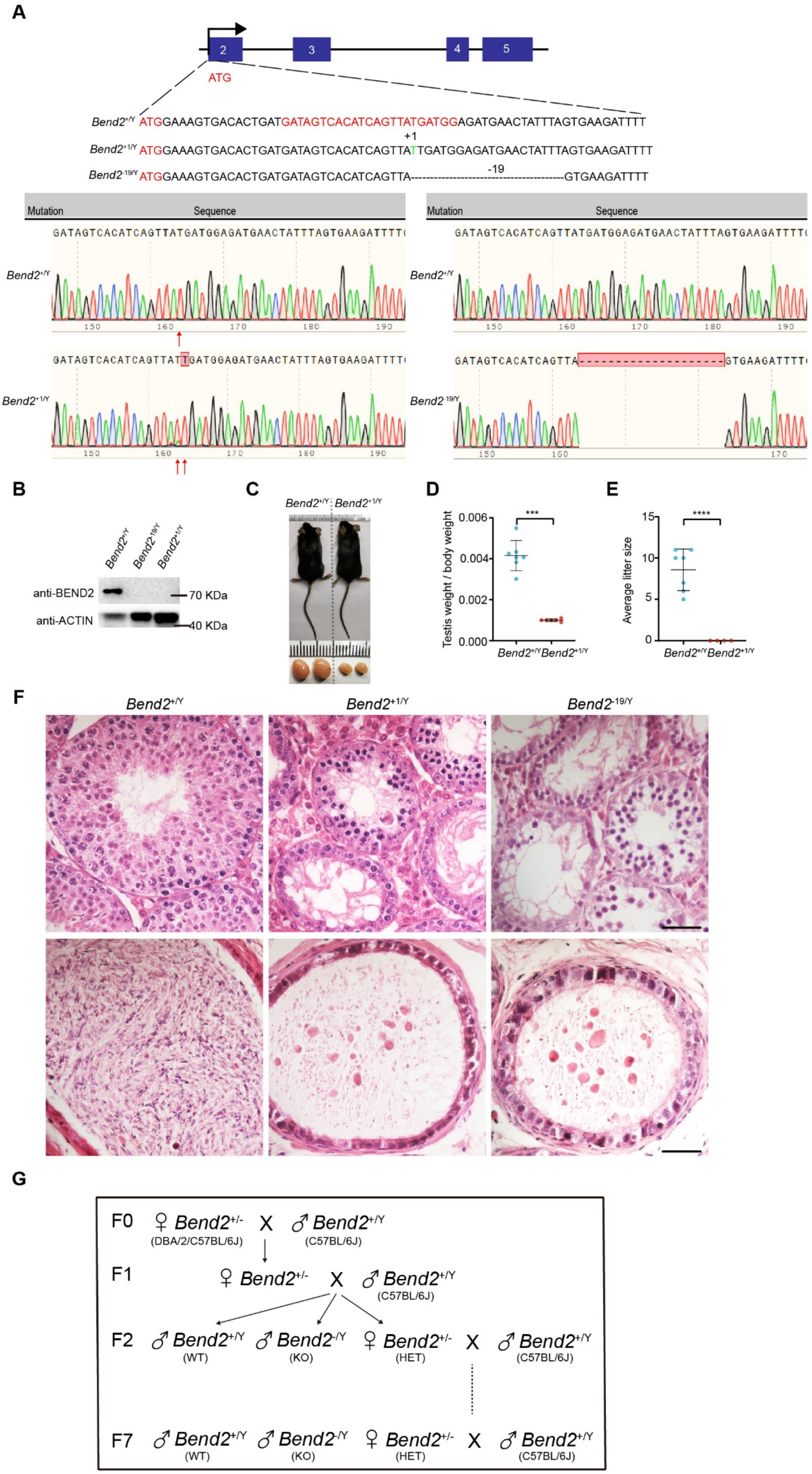
A
2
3
4
5
ATG
Bend2+/Y ATGGAAAGTGACACTGATGATAGTCACATCAGTTATGATGGAGATGAACTATTTAGTGAAGATTTT
+1
Bend2+1/Y ATGGAAAGTGACACTGATGATAGTCACATCAGTTATTGATGGAGATGAACTATTTAGTGAAGATTTT
-19
Bend2-19/Y ATGGAAAGTGACACTGATGATAGTCACATCAGTTA-------------------GTGAAGATTTT
Mutation
Sequence
Bend2+/Y
Bend2+1/Y
Bend2-19/Y
B
anti-BEND2
anti-ACTIN
70 KDa
40 KDa
C
D
Testis weight / body weight
E
Average litter size
F
G
F0 ♀ Bend2+/- X ♂ Bend2+/Y
(DBA/2/C57BL/6J) (C57BL/6J)
F1 ♀ Bend2+/- X ♂ Bend2+/Y
(C57BL/6J)
F2 ♂ Bend2+/Y (WT) ♂ Bend2-/Y (KO) ♀ Bend2+/- (HET) X ♂ Bend2+/Y (C57BL/6J)
F7 ♂ Bend2+/Y (WT) ♂ Bend2-/Y (KO) ♀ Bend2+/- (HET) X ♂ Bend2+/Y (C57BL/6J)

**Fig. S2 *Bend2* is essential for spermatogenesis.**

**(A)** Schematic illustration of the mutant locus in the *Bend2* gene in *Bend2*$^{+1/Y}$ and *Bend2*$^{-19/Y}$ mice, respectively, followed by the alignment of genomic sequence in these two mutant mice. **(B)** Western blot confirmation of the deletion of BEND2 protein both in *Bend2*$^{+1/Y}$ and *Bend2*$^{-19/Y}$ mice using rpAb-B2. **(C)** Note the significant size reduction in 8-week-old *Bend2*$^{+1/Y}$ testes. **(D)** Quantitative comparison of testis/body ratios between *Bend2*$^{+/Y}$ and *Bend2*$^{+1/Y}$ mice (***$p$<0.001, Student's *t*-test). **(E)** Comparison of litter size between *Bend2*$^{+/Y}$ and *Bend2*$^{+1/Y}$ mice (****$p$<0.0001). **(F)** H&E staining of testicular and epididymal sections in *Bend2*$^{+/Y}$, *Bend2*$^{+1/Y}$, and *Bend2*$^{-19/Y}$ mice. **(G)** Schematic diagram of the breeding strategy for *Bend2*$^{-4k/Y}$ mice from founder to the F7 generation.

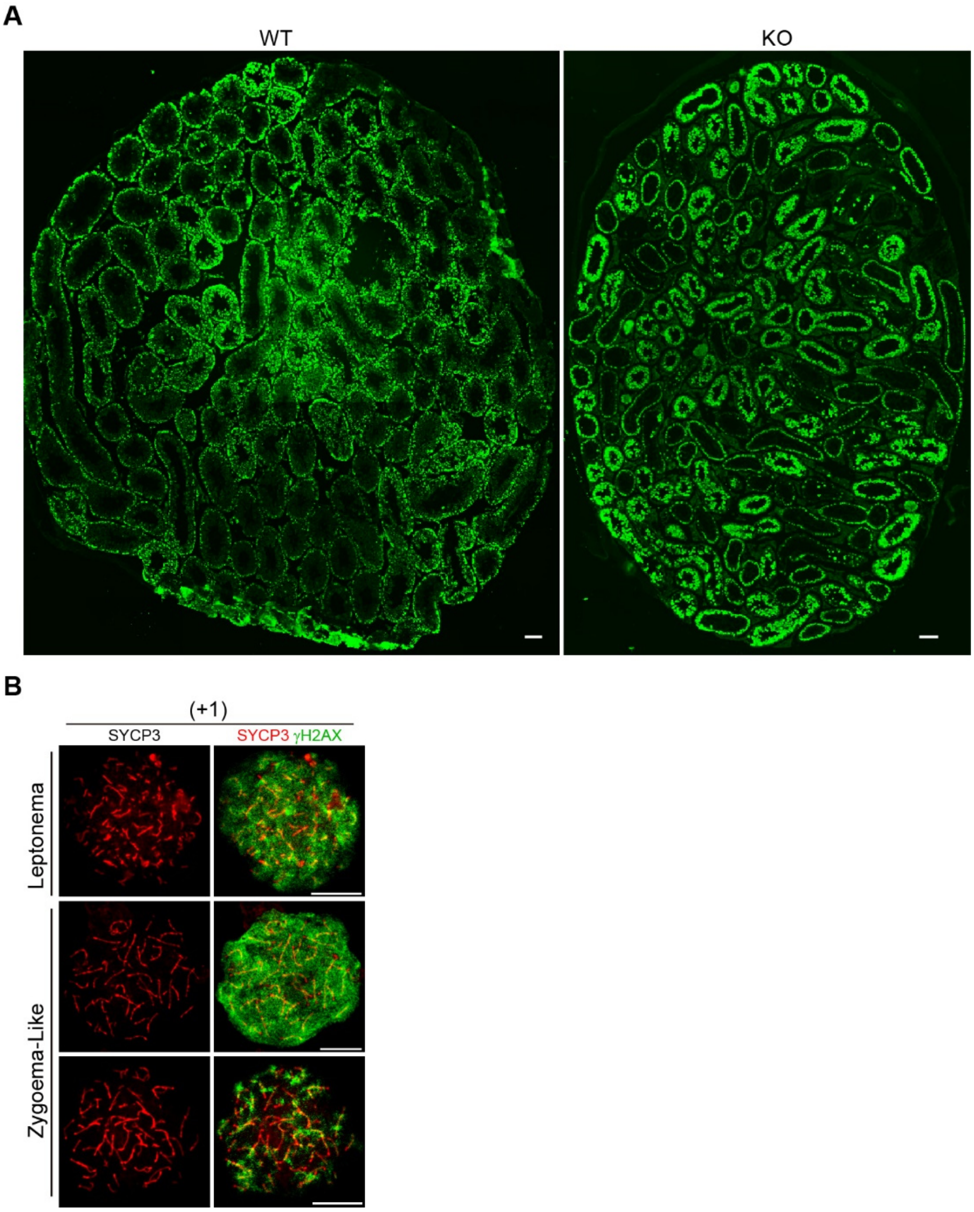


**Fig. S3 Persistence of γH2AX in *Bend2* knockout spermatocytes.**

**(A)** Images of testicular section immunostained with γH2AX antibodies (green) in *Bend2*$^{+/Y}$ and *Bend2*$^{-4k/Y}$ mice (scale bar, 100 μm). **(B)** Nuclear spreads of leptotene and zygotene-like spermatocytes in *Bend2*$^{+1/Y}$ mice. Spermatocytes were immunostained with SYCP3 (red) and γH2AX (green) (scale bar, 10 μm).

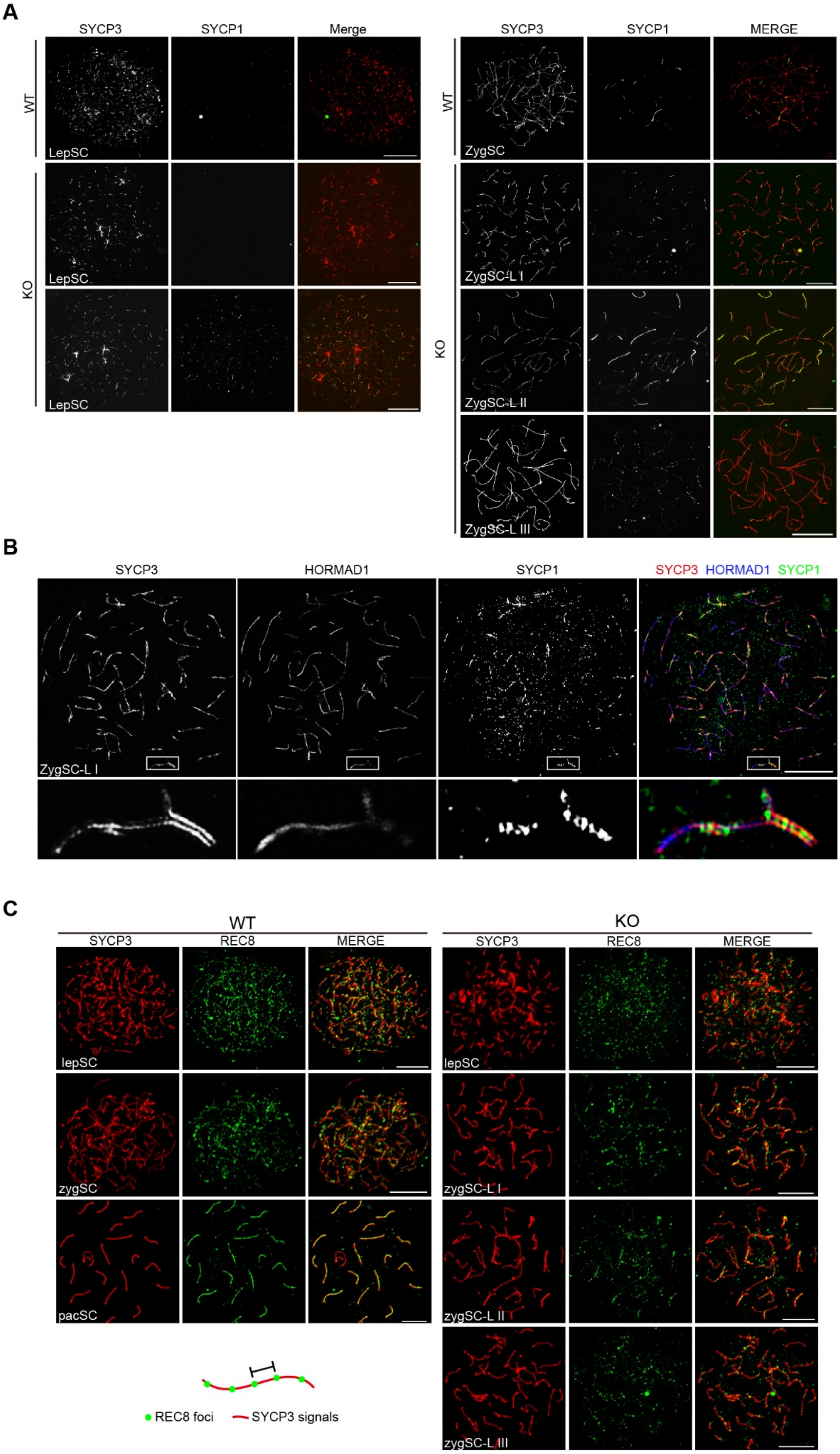
A
SYCP3
SYCP1
Merge
WT
KO
LepSC
LepSC
LepSC
SYCP3
SYCP1
MERGE
WT
KO
ZygSC
ZygSC-L I
ZygSC-L II
ZygSC-L III
B
SYCP3
HORMAD1
SYCP1
SYCP3 HORMAD1 SYCP1
ZygSC-L I
C
WT
SYCP3
REC8
MERGE
lepSC
zygSC
pacSC
KO
SYCP3
REC8
MERGE
lepSC
zygSC-L I
zygSC-L II
zygSC-L III
REC8 foci
SYCP3 signals

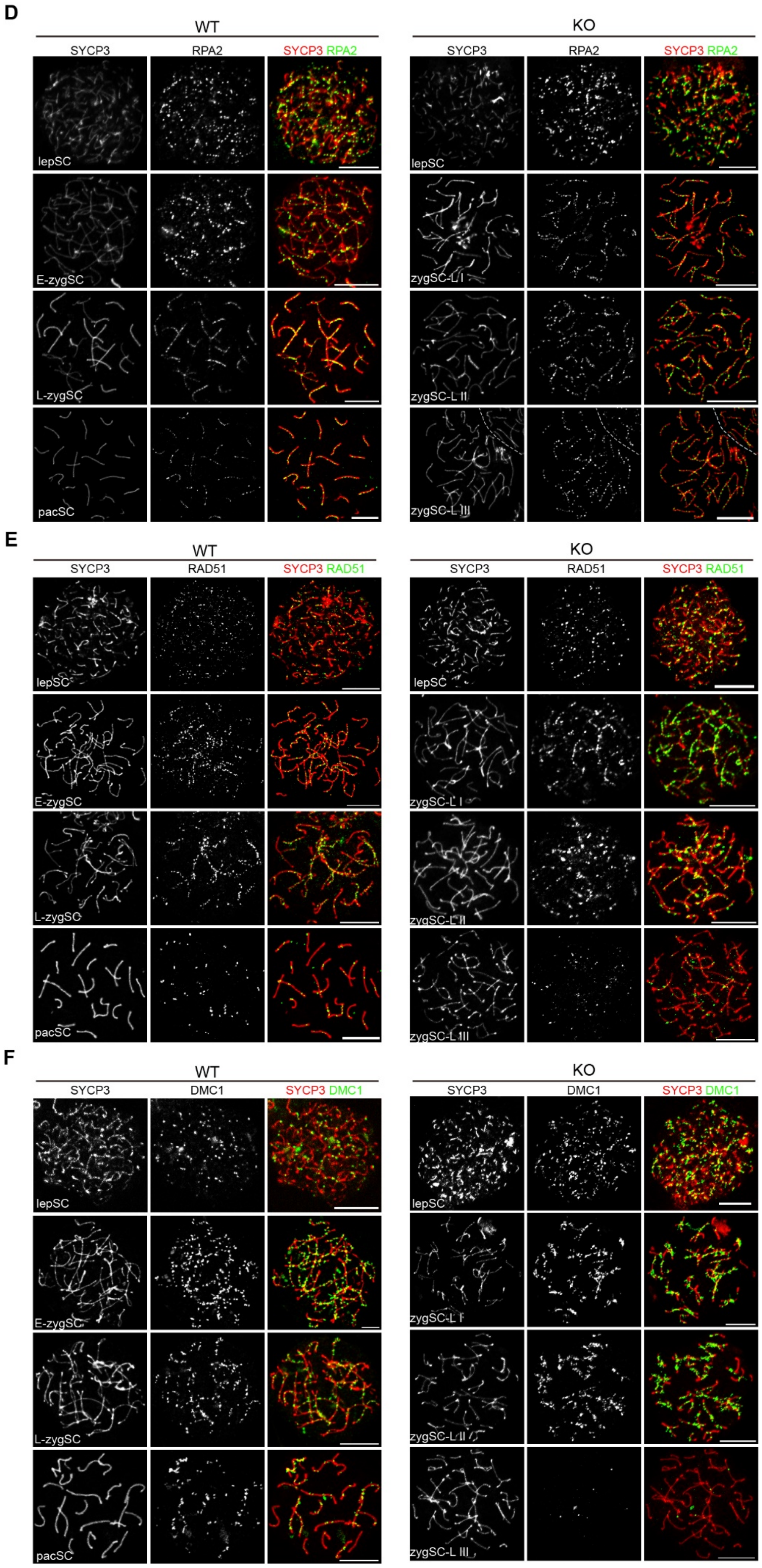

D
WT
KO
SYCP3
RPA2
SYCP3 RPA2
lepSC
E-zygSC
L-zygSC
pacSC
zygSC-L I
zygSC-L II
zygSC-L III
E
RAD51
SYCP3 RAD51
F
DMC1
SYCP3 DMC1

**Fig. S4 BEND2 is essential for chromosomal synapsis in meiosis.**

**(A)** Super-resolution microscopic images of leptotene and zygotene spermatocytes in *Bend2*$^{+/Y}$ and *Bend2*$^{-4k/Y}$ mice. **(B)** Immunofluorescent labeling of SYCP3 (red) HORMAD1 (blue) and SYCP1 (green) in *Bend2*$^{+/Y}$ and *Bend2*$^{-4k/Y}$ mouse spermatocytes. **(C)** Immunofluorescent labeling of SYCP3 (red) and REC8 (green) in *Bend2*$^{+/Y}$ and *Bend2*$^{-4k/Y}$ mouse spermatocytes. **(D)** Immunofluorescent labeling of SYCP3 (red) and RPA2 (green) in *Bend2*$^{+/Y}$ and *Bend2*$^{-4k/Y}$ mouse spermatocytes. **(E)** Immunofluorescent labeling of SYCP3 (red) and RAD51 (green) in *Bend2*$^{+/Y}$ and *Bend2*$^{-4k/Y}$ mouse spermatocytes. **(F)**. Immunofluorescent labeling of SYCP3 (red) and DMC1 (green) in *Bend2*$^{+/Y}$ and *Bend2*$^{-4k/Y}$ mouse spermatocytes (scale bar, 10 μm).

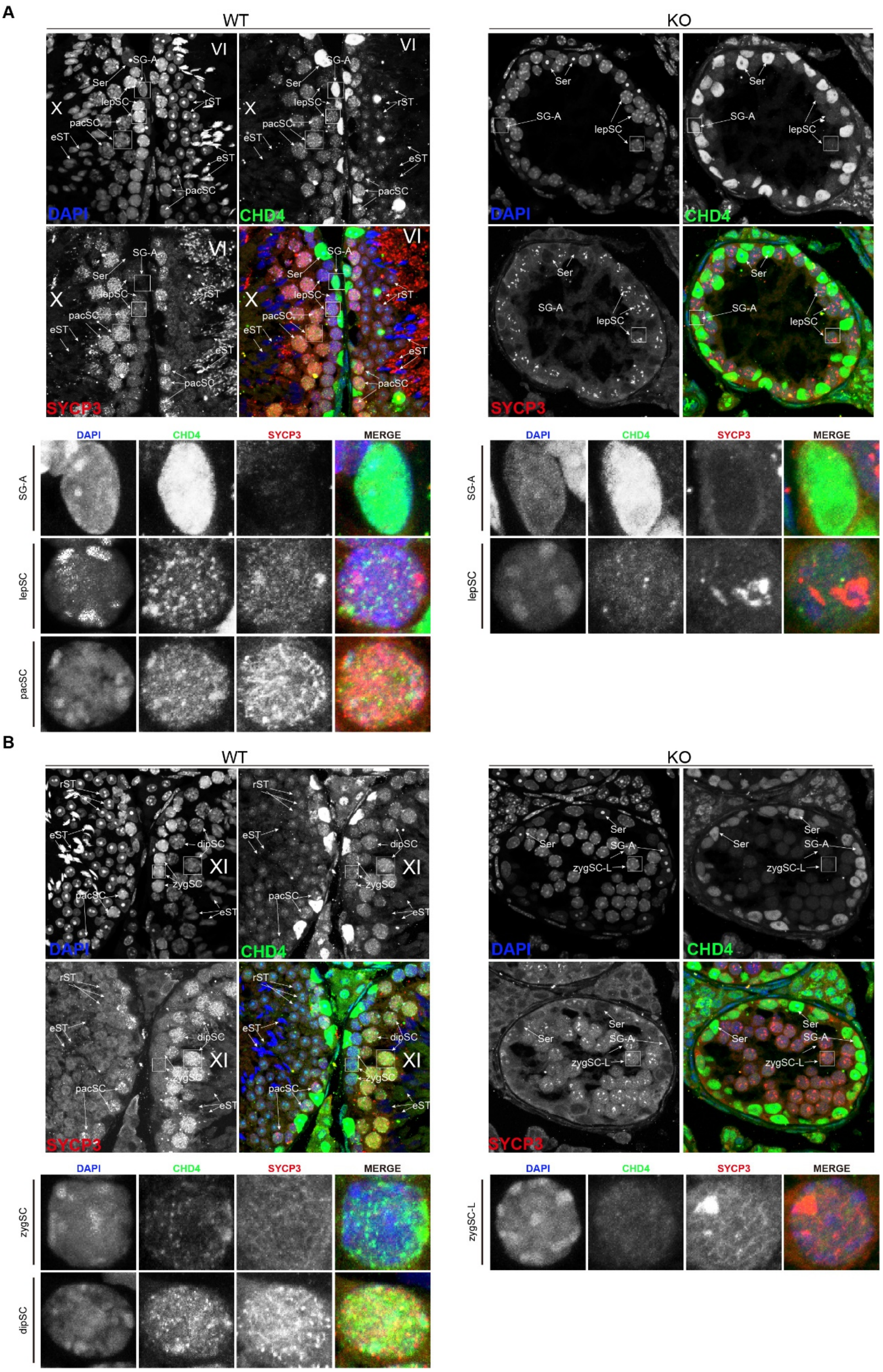


**Fig. S5 The immunofluorescence pattern of CHD4 is altered in *Bend2*$^{-4k/Y}$ mice.**

**(A)** Immunofluorescent staining of SYCP3 (red) and CHD4 (green) in stage VI and stage X sections of WT and KO seminiferous tubules. Enlarged photomicrographs show type A spermatogonia (SG-A), leptotene spermatocytes (lepSC), and pachytene spermatocytes (pacSC). (B) Immunofluorescent staining of SYCP3 (red) and CHD4 (green) in stage XII sections of WT and KO seminiferous tubules. Enlarged photomicrographs show zygotene or zygotene-like spermatocytes (zygSC/zygSC-L) and diplotene spermatocytes (dipSC), respectively.

A

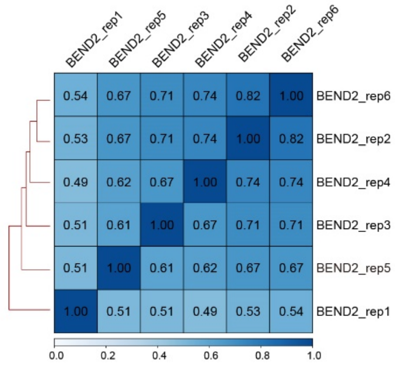


B

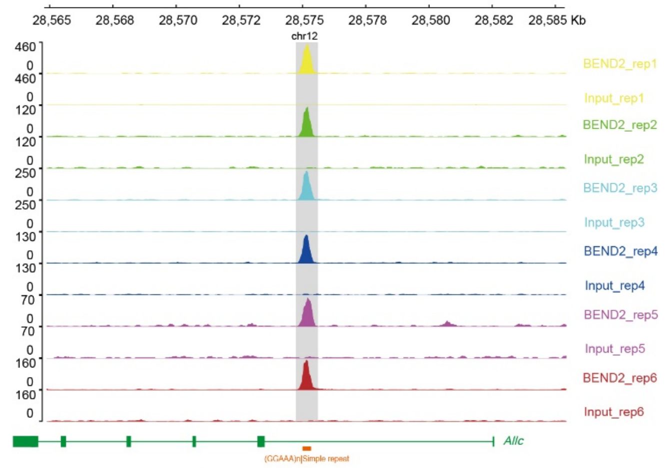


C

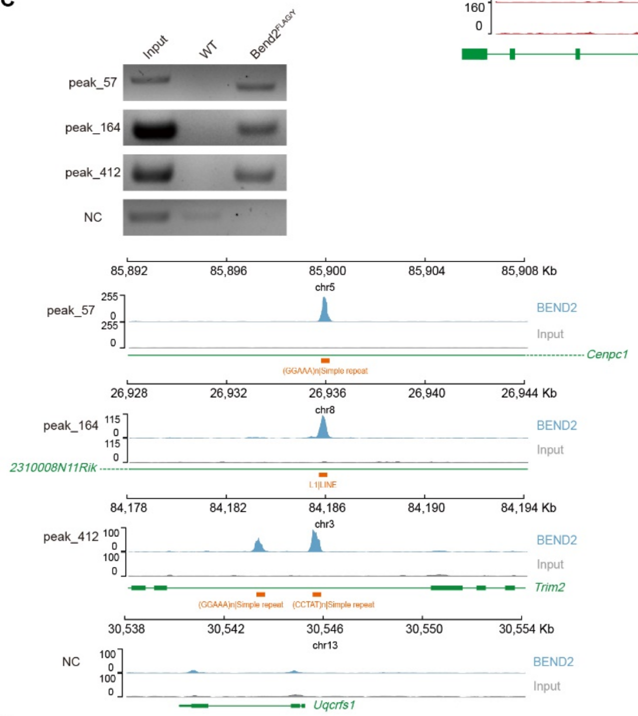


D

| Annotation | Peak(%) | Ratio(obs/exp) | *p* value |
|---|---|---|---|
| 5' UTR | 0.15 | 1.67 | 1.23E-02 |
| Promoter | 1.40 | 1.26 | 3.04E-04 |
| 3' UTR | 0.38 | 0.50 | 1 |
| TTS | 0.95 | 0.94 | 7.75E+01 |
| Exon | 0.64 | 0.50 | 1 |
| Intron | 37.62 | 1.08 | 1.16E-13 |
| Intergenic | 58.72 | 0.97 | 1 |
| other | 0.15 | 0.68 | 1 |

| Annotation | Peak(%) | Ratio(obs/exp) | *p* value |
|---|---|---|---|
| CpG-Island | 0.25 | 2.17 | 2.91E-06 |

| Annotation | Peak(%) | Ratio(obs/exp) | *p* value |
|---|---|---|---|
| Non-repeat | 31 | 0.55 | 1 |
| Repeat | 69 | 1.58 | < 2.22E-16 |

| Annotation | Peak(%) | Ratio(obs/exp) | *p* value |
|---|---|---|---|
| LINE | 11.07 | 0.53 | 1 |
| SINE | 5.32 | 0.71 | 1 |
| Low_complexity | 8.21 | 11.02 | < 2.22E-16 |
| LTR | 9.07 | 0.75 | 1 |
| Simple_repeat | 35.02 | 15.84 | < 2.22E-16 |
| Satellite | 0.31 | 1.76 | 1.83E-04 |

E

Table of significant motifs in BEND2 peaks

| Rank | Motif | P-value | % of Targets | % of Background |
|---|---|---|---|---|
| 1 | AGGACAGGACAG | 1e-5036 | 44.38% | 4.63% |
| 2 | AAAGGAAAAG | 1e-2673 | 24.90% | 2.54% |
| 3 | GGATGGATGGAT | 1e-1592 | 10.29% | 0.48% |
| 4 | CCTAACCATACC | 1e-903 | 9.97% | 1.22% |
| 5 | ATATCTCCCTTG | 1e-484 | 2.28% | 0.05% |
| 6 | GGTAGGTA | 1e-406 | 21.05% | 9.75% |
| 7 | TCTATCATCTAT | 1e-381 | 4.24% | 0.51% |
| 8 | GGGCCTCTCACT | 1e-315 | 1.50% | 0.03% |

Table of *de novo* motifs

| Rank | Motif | P-value | % of Targets | % of Background | Best Match |
|---|---|---|---|---|---|
| 1 | AAGGAAAGGAAA | 1e-3637 | 41.49% | 1.49% | UME1<br>PCBP1 |
| 2 | GAAAAGAAAAGA | 1e-1092 | 24.98% | 3.20% | AEF1 |
| 3 | AAGGGTAGGG | 1e-835 | 30.27% | 6.89% | HRB98DE |

Table of known motifs

| Rank | Motif | Name | P-value | % of Targets | % of Background |
|---|---|---|---|---|---|
| 1 | CGGAAGTGAAAC | PU.1 | 1e-1578 | 81.23% | 34.25% |
| 2 | TGGAACAGAA | ZNF189 | 1e-414 | 34.60% | 14.73% |
| 3 | TGGGGAAGGGCA | ZNF467 | 1e-358 | 33.26% | 14.85% |

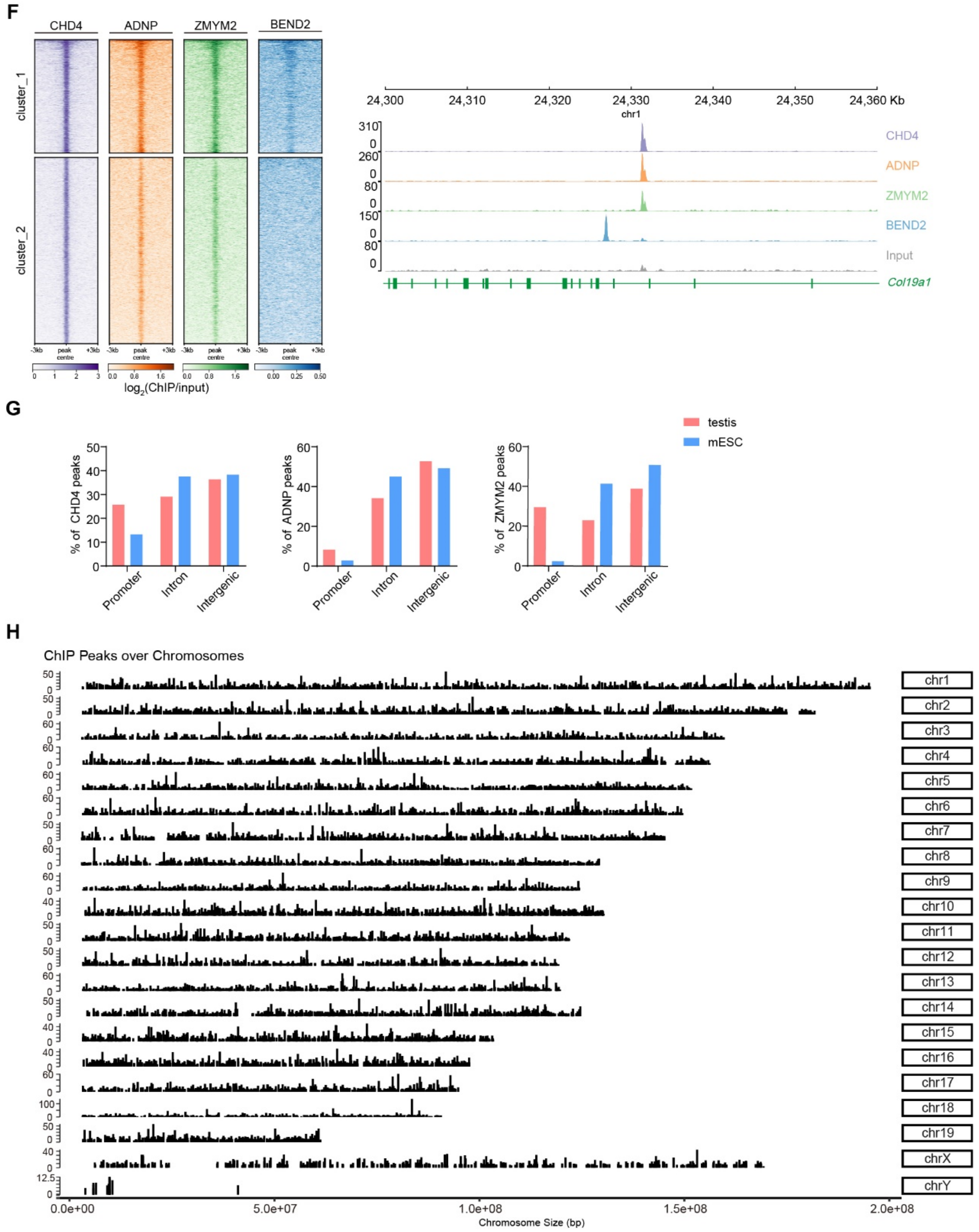


**Fig. S6 Characteristic analysis of the BEND2-binding site.**

**(A)** Heatmap analysis showing the correlations among six BEND2 ChIP-seq replicates. **(B)** Browser view showing ChIP-seq signals of BEND2 (six biological replicates, rep1-6). **(C)** ChIP-PCR analyses and browser view of BEND2-binding sites. **(D)** A short table comprising the annotation of BEND2 DNA-binding sites (up). Repeat analysis of BEND2-targeted sites in introns and intergenic regions (down). **(E)** BEND2 DNA-binding motifs predicted by HOMER; *p*-values and frequencies of occurrence for motif enrichment compared to genomic background are indicated. **(F)** Heatmap of CHD4, ADNP, ZMYM2, and BEND2 ChIP-seq enrichment across all CHD4 peak midpoints, with each row representing a 6-kb window centered on CHD4 peak midpoints (left). Cluster_1 represents CHD4, ADNP, ZMYM2, and BEND2 enriched in a sub-class of CHD4 DNA-binding sites. Cluster_2 represents CHD4, ADNP, and ZMYM2 enriched in a sub-class of CHD4 DNA-binding sites. An example of CHD4, ADNP, and ZMYM2 co-binding site without enrichment of BEND2 (right). **(G)** Genome feature-distribution analysis of CHD4, ADNP, and ZMYM2 peaks in testis and ESC. **(H)** Chromosomal distribution of BEND2 ChIP-seq peaks.

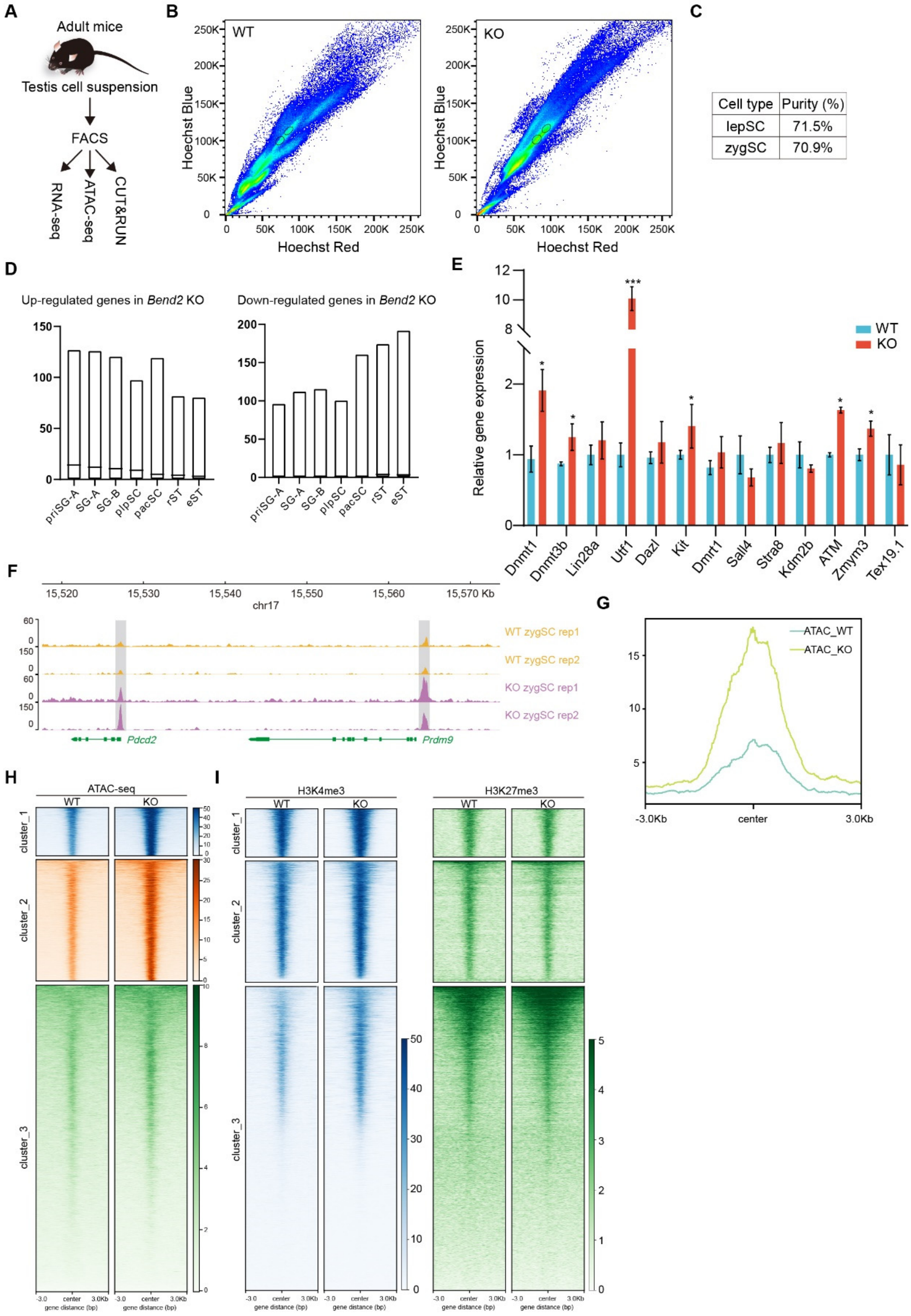


**Fig. S7 Transcriptome analysis of spermatocytes in meiotic prophase I from *Bend2* KO mice.**

**(A)** Experimental workflow. **(B)** Spermatocytes were stained with Hoechst 33342 and sorted by FACS to enrich for either leptonema- or zygonema-stage cells. **(C)** The cell purity of leptonema- or zygonema-stage cells as sorted by FACS; SYCP3 and γH2AX were used to identify cell stage. **(D)** The upregulated and downregulated genes in *Bend2* KOs were re-analyzed vis-à-vis previously published data of stage-specific bulk RNA-seq (N = two biologically independent samples (*52*)]). **(E)** Validation of differentially expressed genes using real-time PCR analysis (Student's *t*-test: *, $p<0.05$; ***, $p<0.001$). **(F)** Browser view showing ATAC-seq signals of spermatocytes at zygonema (N = two biologically independent samples). **(G)** The distribution of ATAC-seq signal around the TSS region, signifying the BEND2-bound promoter genes. **(H)** Heatmap of ATAC-seq enrichment across all TSSs, with each raw datum representing a 6-kb window centered on TSS midpoints. **(I)** Heatmap of H3K4me3 and H3K27me3 levels on the TSS of three clusters of genes.
